# Behavioral Plasticity and Dispersal of Free Swimming Coral Larvae via Vertical Tracking Microscopy

**DOI:** 10.64898/2026.11.19.767947

**Authors:** Ian Ho, Qing Zhang, Benjamin Foster, Elora H. López-Nandam, Pranav Vyas, Shayle Matsuda, Rebecca Albright, Manu Prakash

## Abstract

Coral larvae disperse and seek suitable sites for settlement while undergoing rapid development, making this period central to reef connectivity, recruitment, and recovery. Yet how the accompanying changes in larval morphology, ciliary activity, and swimming behavior shape dispersal remains poorly understood. Here, using vertical tracking microscopy, we follow individual *Acropora mille-pora* larvae over hundreds of body lengths at successive developmental stages while resolving body morphology. We show a developmental progression from buoyant rise to upward- and downward-directed gravitactic swimming, followed by weaker gravitactic alignment at later stages, accompanied by body elongation and increasingly structured ciliary flows. Integrating these kinematics into a minimal, data-driven model, we separate buoyant drift from active swimming and estimate the propulsion costs of vertical migration and lateral spreading. Buoyancy supports early ascent without propulsion expenditure, while the model predicts lower costs of descent and lateral spreading at late developmental stages. Simulations in fluctuating flow, assuming no flow sensing, predict that directed swimming weakens when flow-induced rotation acts on timescales comparable to intrinsic heading relaxation. Together, these results reveal how development reorganizes the physical drivers of larval motion in ways that may support the transition from dispersal to settlement, providing a basis for incorporating developmental changes in swimming into models of coral larval dispersal.

## I. INTRODUCTION

Coral colonies are fixed in place, but their populations are connected by the journeys of larvae through the ocean. The propagation, dispersal, and settlement of these larvae govern the connectivity of coral populations across reef systems and ultimately shape the distribution, genetic structure, and resilience of reef communities [1]. Corals occupy a diverse range of marine environments, spanning shallow tropical reefs, mesophotic reefs, and deep-sea ecosystems, forcing larvae to navigate distances of O(10^3^)–O(10^6^) body lengths on their way to settlement [2–5]. Larval replenishment has become increasingly important as reefs face increasing stressors, including strong storms, predator outbreaks, and climate-driven coral bleaching events [6]. Supply of larvae from other reefs and settlement at favorable sites both strongly improve reef recovery [7–10]. An understanding of larval dispersal and settlement is therefore central to both fundamental questions of marine population connectivity [11] and to forecasting reef trajectories under mounting environmental pressure. Dispersal reflects the interplay of larval swimming dynamics, and the local hydrodynamic environment [12].

The transition from dispersal to settlement places changing demands on larval motion over the course of its life cycle (Fig. 1a). In many broadcast-spawning corals, buoyant egg–sperm bundles rise toward the surface, where increased encounters among gametes promote fertilization [13]. Embryos and larvae then undergo a period of planktonic development and dispersal. As larvae become competent to settle, they can descend toward the benthos and explore potential settlement sites before attachment and metamorphosis [14, 15]. Vertical motion throughout the planktonic phase reflects a changing balance between buoyancy and active swimming [16]. Gravitactic swimming has been reported across several coral taxa [17–19], including steady ascent or descent with occasional reversals in direction [17]. This motility is powered by coordinated ciliary beating, and its effectiveness is influenced by larval morphology, age, and physiological state [20, 21]. Moreover, swimming speed and preferred vertical direction vary among species and developmental stages [20, 22–25].

**FIG. 1.**
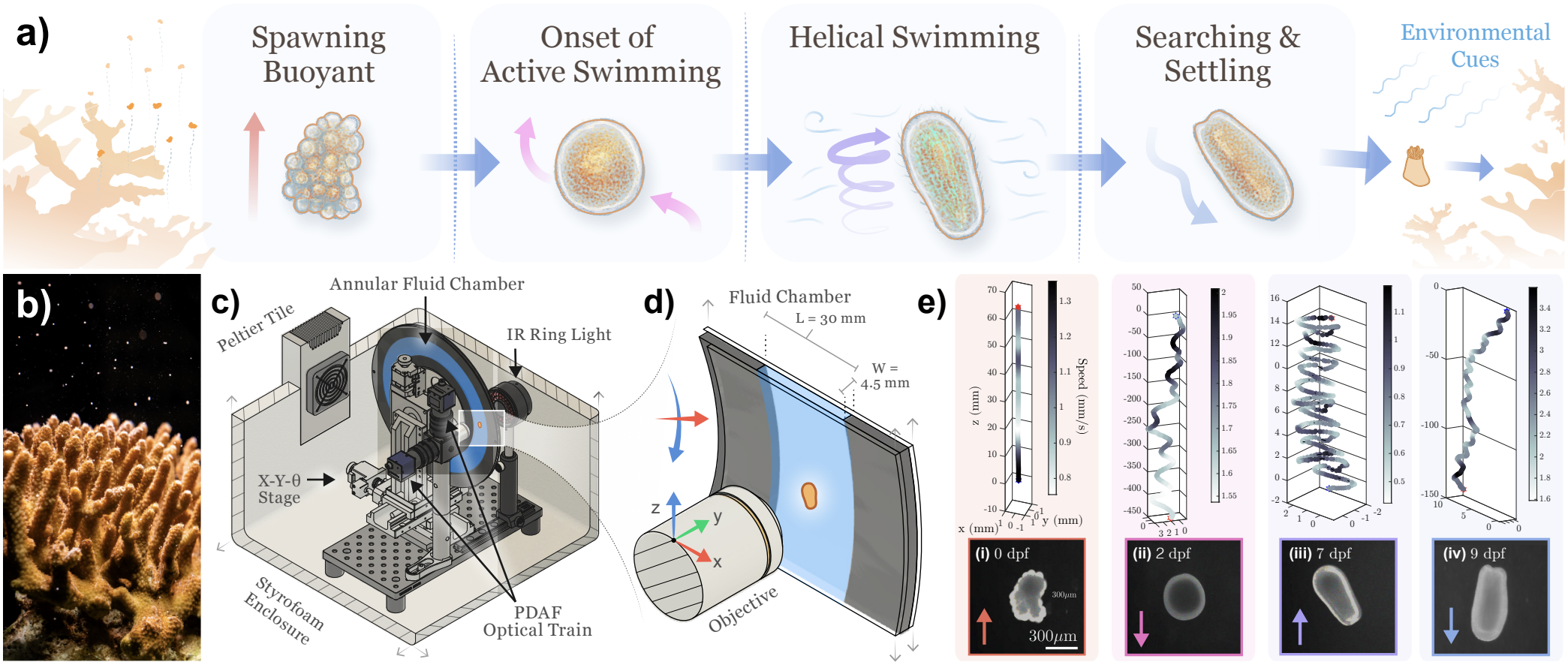
Vertical tracking microscopy reveals developmental changes in Acropora millepora larval behavior. Schematic of the coral life cycle from broadcast spawning to free-swimming larval stages and, ultimately, settlement and metamorphosis. Across this interval, larvae transition from predominantly buoyancy-driven motion at the earliest stages to increasingly coordinated swimming at later stages. (b) *Acropora millepora* spawning. (c) Schematic of the vertical tracking microscope housed in a temperature-controlled enclosure. An annular fluid chamber mounted on *x*− *y*−*θ* stages allows extended motion along the gravity axis while maintaining the larva within the microscope field of view. (d) Infrared dark-field imaging and closed-loop feedback are used to compensate larval motion by stage translation (*x*) and chamber rotation (*z*); focal position along (*y*) is adjusted manually. Illustrated fluid chamber and larvae are not to scale. (e) Representative trajectories across development, characterized by days post fertilization (dpf). Immediately after cleavage, motion is dominated by buoyancy; by ∼ 2 dpf larvae exhibit downward bouts against buoyant rise; by ∼ 7 dpf many larvae show coordinated helical swimming; and by ∼ 9 dpf trajectories exhibit substantial lateral excursion. Photo credit for panel (b): Gayle Laird (California Academy of Sciences).

Larval motion is shaped by both external sensory cues and internal developmental state. Coral larvae can respond to light, chemical cues, and sound, which can influence their swimming behavior and selection of settlement sites [26–30]. Responses to light may help larvae locate settlement habitats suitable for the photosynthetic requirements of the settled coral, even in species whose larvae initially lack algal symbionts [31, 32]. Chemical cues associated with crustose coralline algae can also influence larval selection of settlement substrates [29, 33], while epidermal cilia have been proposed to detect the oscillatory water motion associated with sound [34]. Mechanical disturbance has also been associated with changes in coral larval swimming, including reduced swimming activity following water agitation in larvae of *Eunicella singularis* [35]. Physiological changes also shape larval motion. Maternally supplied lipid reserves support development and propulsion [36, 37] while contributing to positive buoyancy [38]. Their abundance and composition change as larvae mature, and depletion of these reserves can increase larval density and reduce buoyant support [39, 40]. Environmental cues and development therefore shape larval motion through distinct routes: cues modulate active behavior, whereas development alters both active swimming and passive buoyancy.

Here, we focus on how swimming and buoyancy change across development, rather than on responses to specific environmental cues. Motility develops alongside changes in settlement competence and sensory organization. Larval precompetency and settlement behavior varies widely between species, with a reported precompetency period of approximately 2–6 days for the Indo-Pacific broadcast-spawning species [41]. Survival and settlement competence also vary with age. In *Acropora millepora*, the focused species of our study, the modeled proportion of larvae both alive and competent to settle has been reported to peak approximately 8–10 days after spawning [42]. Over this interval, body form and neural organization develop, and changes in neuropeptide expression accompany the progression toward settlement and metamorphosis [43]. These concurrent changes place swimming behavior within a rapidly evolving developmental context.

However, how these developmental changes affect dispersal also depends on the surrounding flow. Larvae encounter a wide range of flow scales and flow structures as they disperse and settle, from reef turbulence [44] to topographically generated [45] and buoyancy-driven fronts advected by wind and local currents [46–48]. Active swimming allows larvae of many marine species to vertically migrate across depth-varying flow layers, potentially enhancing or mitigating environmental advection depending on their behavioral strategy [49–52]. Near-surface transport can additionally favor dispersal away from reefs, with simulations of Great Barrier Reef circulation predicting faster dispersal and reduced local retention under surface currents than under depth-averaged flow [53]. Vertical position in the water-column therefore modulates dispersal trajectories, residence time near suitable substrates, and exposure to predation or unfavorable conditions. By contrast, flow gradients can also reorient larvae, making heading dynamics important to the persistence of directed transport [49]. Models of individual coral larvae show how swimming and flow-induced rotation interact with substrate topography to influence settlement [47, 54], but use fixed swimming speeds and body shapes without resolving developmental changes in buoyancy or intrinsic heading dynamics. Accurately representing larval behavior remains a central challenge in biophysical models of dispersal [55].

We hypothesize that developmental changes in buoyancy and swimming help larvae meet the contrasting demands of dispersal and settlement while drawing on limited maternal energy reserves. Early buoyant ascent requires no propulsion expenditure or maintained swimming heading, so its upward contribution persists when flow reorients the larva. Active swimming subsequently permits descent against residual positive buoyancy, while depletion of lipid reserves can reduce that buoyancy and make descent less costly. Selection may therefore shape the timing of both buoyancy changes and swimming behaviors across development, linking the physical drivers of larval motion to the changing demands of the life cycle.

Previous work has documented ontogenetic changes in vertical swimming behavior across multiple coral species [16, 20, 22, 56]. Yet how these changes interact with buoyancy to shape vertical transport, lateral spreading, and how the energetic demands of swimming vary across development, remains poorly understood. To connect microscale behavior with transport over longer distances, we use vertical tracking microscopy to follow individual *A. millepora* larvae over hundreds of body lengths at successive developmental stages through 9 days post fertilization, while resolving body morphology. Complementary ciliary-flow measurements and neural staining place these swimming states in their developmental context. We combine the trajectories with geometry-based estimates of buoyant drift for individual larvae to constrain a minimal model that separates buoyancy, directed swimming, and stochastic reorientation. This integration allows us to estimate how the propulsion costs of vertical migration and lateral spreading change across development. Simulations in fluctuating flow then assess how much influence these behaviors retain on transport when larvae respond to gravity alone, providing a baseline without flow sensing.

## II. EXPERIMENTAL METHODS

In this study, adult *Acropora millepora* colonies were maintained and spawned *ex situ* at the California Academy of Sciences Coral Regeneration Lab (Fig. 1b). Spawning was induced by reproducing seasonal temperature and lunar light cycles in a light-controlled aquarium system. Here, we tracked *A. millepora* larvae longitudinally across development, from immediately after fertilization through 9 days post fertilization, spanning the sequence of early buoyant stages, the onset of active swimming, and late pre-settlement behavior.

Larvae were imaged in a vertical tracking microscope based on the hydrodynamic-treadmill architecture of Krishnamurthy *et al* [59]. A vertically oriented annular chamber was mounted on *x*–*y*–*θ* stages and used with closed-loop feedback to compensate organism motion by chamber rotation and stage translation while maintaining microscopic resolution over extended virtual vertical distances (Fig. 1c,d). A temperature-stabilized chamber, housed within a styrofoam enclosure and regulated with Peltier elements, maintained the experimental temperature at 26^*°*^C (Fig. 1c). Larvae were tracked in a light-tight enclosure using infrared back-illumination to minimize perturbation by visible-light cues during long-duration recordings (Fig. 1d). This precaution was important because abrupt light dimming in *A. millepora* can induce whole-body contraction, ciliary arrest, and a marked reduction in swimming speed [60]. Images were acquired at 15–50 fps using DaHeng USB cameras (MER-160-227U3M) at 4× and 10× magnification. Larval position in the field of view was extracted in real time using *DaSIamRPN*, a distraction aware Siamese network [61], and the resulting tracking error was used to drive compensatory *x*-translation and *θ*-rotation to keep the larva mostly centered in the field of view (Fig. 1d).

The imaging chamber is an annulus of inner radius *R*_*i*_ = 80 mm, outer radius *R*_*o*_ = 110 mm, radial thickness *L* = *R*_*o*_ − *R*_*i*_ = 30 mm, and chamber width *W* = 4.5 mm. Across all tracked larvae, the body major- and minor-axis dimensions were *ℓ*_*∥*_ = 0.87 ± 0.23 mm and *ℓ*_*⊥*_ = 0.42 ± 0.08 mm, respectively (mean ±s.d., *n* = 345 trajectory-level measurements). Because larvae were predominantly gravitactic and generally aligned with their long axis along the effective vertical direction, confinement relative to the chamber cross-section can be quantified using the minor-axis body dimension, giving *ℓ*_*⊥*_*/W* = 0.093 ±0.017 and *ℓ*_*⊥*_*/L* = 0.0140 ±0.0026. Focal position along *y* was adjusted manually during experiments, and we therefore use the projected *x*–*z* coordinates as the most reliable representation of larval kinematics in this setup.

Before each experiment, the annular chamber was filled with artificial seawater with density ≈ 1.028 g cm^*−*3^ and equilibrated within the temperature-controlled enclosure. In selected experiments, the seawater was seeded with 1*µ*m-diameter tracer particles for flow visualization or particle image velocimetry (PIV). Larvae were then introduced into the chamber at a density of approximately 15–20 individuals per wheel (fluid volume ≈ 70 mL), which provided enough animals for efficient sampling while reducing the likelihood of repeatedly tracking the same larva. Individuals were selected for tracking at random, and each track was typically maintained for approximately 30 s to 2 min, with the goal of sampling as many distinct larvae as possible within each day. Recording durations varied because tracking ended when the swimmer was lost, encountered a chamber boundary, or acquisition was stopped to follow another individual. Although repeat sampling of the same larva cannot be ruled out entirely, the loading density was chosen to reduce this possibility, and individual trajectories were therefore treated as biological replicates in the analysis. For analysis, trajectories were grouped by nominal days post fertilization (dpf), with recording dates, acquisition windows, and sample counts reported in SM, Sec. II.A, Table S1. Subsequently, swimming metrics were weighted by inter-frame duration and normalized within each trajectory so that each trajectory contributed unit mass to the day- level distributions, thereby avoiding over-representation of long tracks. At later developmental stages, when larvae exhibited larger lateral excursions, wall approaches became more frequent. To reduce boundary effects, trajectories were truncated in post-processing once a larva approached within three body lengths (∼ 3*ℓ*_*∥*_) of a radial sidewall or within one body length (∼ *ℓ*_*∥*_) of a front/back wall.

## III. RESULTS

### A. Developmental progression from buoyant motion to active swimming

Across development, larvae pass through a sequence of qualitatively distinct behavioral regimes accompanied by coupled changes in body form and ciliary flow generation (Fig. 1e, 2a–d). Early post-fertilization stages are rounded (Fig. 2a,b) and their motion is dominated by buoyancy, yielding trajectories that are largely monotonic along the gravity axis and showing no clear evidence of active swimming (Fig. 2d, columns 1–2). Consistent with previous reports of mucus secretion in coral embryos and larvae [21, 62], we observed apparent mucus trails behind rising embryos (Fig. 2c,i; SM Movie S1). These trails could modify hydrodynamic drag and hence the rate of buoyant rise. At 1 dpf, we observed the first signs of cilia-driven flow (SM Movie S1), but the trajectory data did not yet indicate propulsion strong enough to appreciably alter larval motion away from its vertical trajectory (Fig. 2d). At approximately 2 dpf (Fig. 2d), larvae first exhibit sustained downward bouts, marking the onset of active propulsion sufficient to oppose buoyant rise. Over the following days, the body becomes more elongated (Fig. 2a,b), the surrounding flow fields become structured (Fig. 2c). FMRFamide immunostaining concurrently reveals a developing ectodermal nerve net (Fig. 2c)[43, 63], which may provide a substrate for sensory modulation of ciliary propulsion and whole-body swimming [60]. Trajectories are highly aligned with gravity, and we observe clear helical swimming in many larvae (Fig. 1e, iii ; SM Movie S1). By 8–9 dpf, this strongly gravitactic mode weakens and the trajectories show larger lateral excursions, horizontal bouts, and occasional reversals or U-turns between upward and downward swimming.

**FIG. 2.**
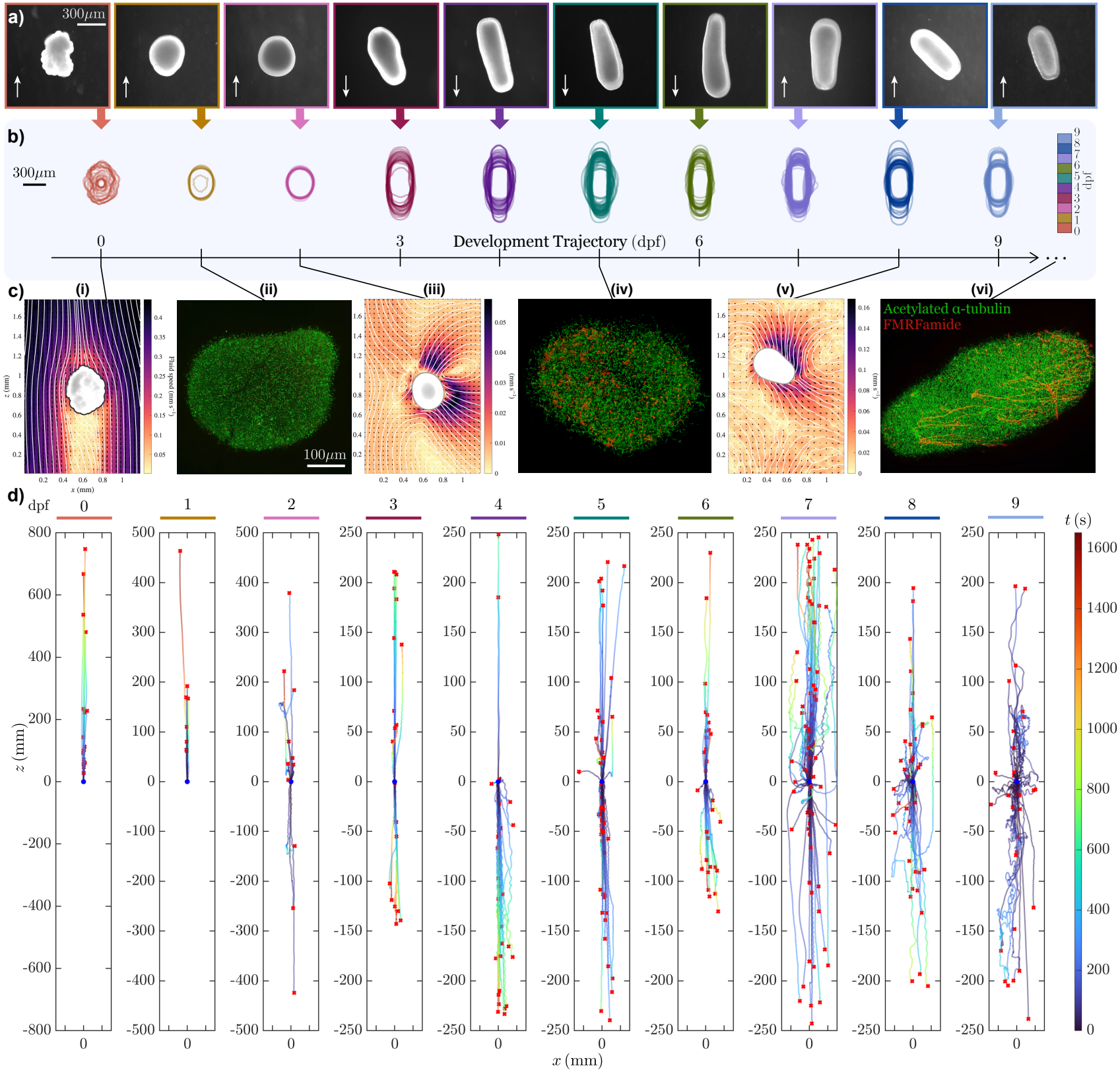
Ontogenetic changes in morphology, ciliary flows, and swimming trajectories. (a) Representative images spanning larval development. Arrows indicate the swimming direction, typically with the broader aboral pole leading. Superposed body outlines show the progression from rounded early stages to elongated planulae. (c) Representative particle-image-velocimetry fields [57] (i, iii, v) and immunostaining images (ii, iv, vi) at selected stages reveal concurrent changes in ciliary forcing and neural development. PIV velocities are shown in a frame translating with the embryo in (i) and in the laboratory frame in (iii, v) at 0,2,8 dpf respectively. Immunostaining images are confocal maximum-intensity projections showing acetylated *α*-tubulin (green, highlighting cilia) and FMRFamide immunoreactivity (red, highlighting a subset of neurons) at 1,5,12 dpf respectively. See Supplemental Material (SM, Sec. II.D.) for staining methods [58]. (d) Projected (*x* − *z*) trajectories for the same developmental sequence (0–9 dpf, from left to right). Trajectories span different observation durations. These trajectories are shown in two dimensions because focal position along (*y*) was adjusted manually during tracking; the (*x* − *z*) projection therefore provides the most consistent kinematic comparison across stages.

### B. Kinematic statistics resolve successive developmental swimming states

Having established the qualitative progression of larval motion, we quantify the swimming trajectories by reporting four complementary kinematic features: vertical swimming velocity, lateral swimming velocity, preferred directions, and directional persistence (Fig. 3). As expected, at 0–1 dpf, the vertical-velocity distributions are narrow and strongly biased in the direction of buoyant rise, while horizontal velocities remain concentrated near zero (Fig. 3a,b). The appearance of negative vertical velocities at 2 dpf marks the onset of active swimming capable of opposing buoyancy. At subsequent stages, the vertical distributions become multi-modal, developing distinct upward- and downward-moving peaks, whereas the horizontal distributions remain centered near zero but broaden, particularly at late stages. At the statistical level, larvae therefore increase their lateral motion without developing a systematic horizontal bias across the sampled trajectories.

**FIG. 3.**
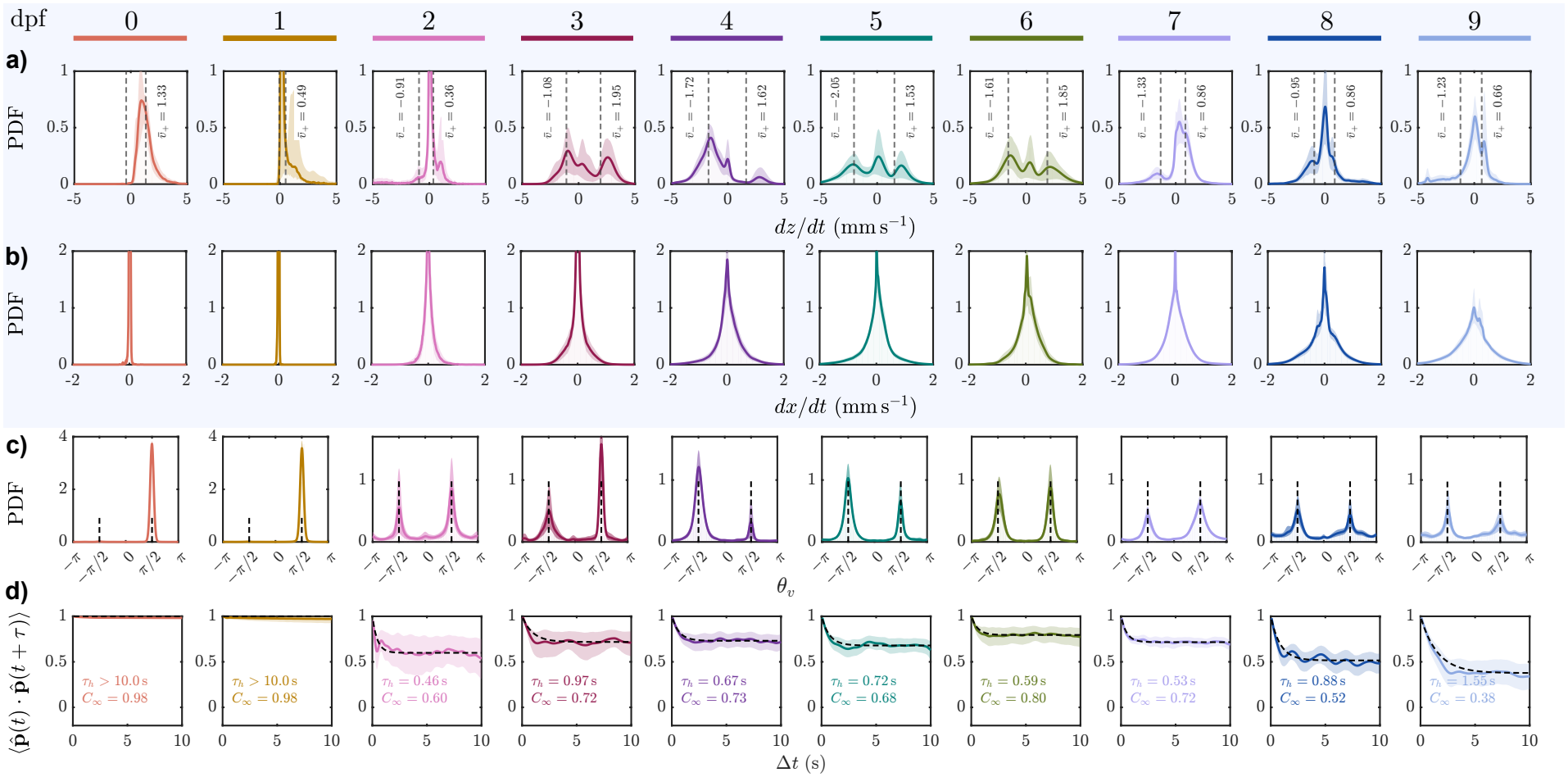
Stage-dependent statistics resolve successive larval swimming states. (a) Probability densities of vertical velocity *ż* across development. Distinct upward- and downward-moving branches emerge after the onset of active swimming; their mean velocities are indicated. (b) Horizontal-velocity densities remain centered near zero but broaden at later stages, indicating increased lateral motion without persistent horizontal drift. (c) Probability densities *π*(*θ*_*v*_) of the projected direction of motion 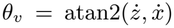. Active stages develop upward- and downward-directed states centered near *θ*_*v*_ = ±*π/*2. (d) Trajectory-heading autocorrelation 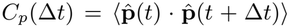. Fits yield a short-time heading-persistence scale *τ*_*h*_ and a long-time plateau *C*_*∞*_. The dpf 0–1 values of *τ*_*h*_ exceed the 10-s fitting window and reflect monotonic buoyant rise rather than active orientation control. Shaded regions denote 95% bootstrap confidence intervals.

To quantify the preferred directions of motion, we define a 2D heading due to the combined action of active swimming and buoyancy as

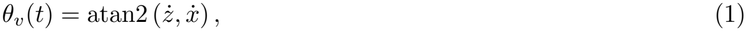

where atan2 is the four-quadrant inverse tangent, and *θ*_*v*_ = *π/*2 or *θ*_*v*_ = −*π/*2 correspond to either upward- or downward-directed motion respectively. Consistent with the vertical-velocity distributions, the heading distributions evolve from a single buoyancy-dominated direction at the earliest stages to distinct upward- and downward-directed states after active swimming begins (Fig. 3c). Likewise, at 8–9 dpf, increased probability weight at oblique and horizontal headings accompanies the broadening of the horizontal-velocity distributions. The heading distributions provide a particularly direct representation of directional bias in *A. millepora* swimming, offering strong evidence for gravitaxis consistent with previous observations in *A. tenuis* [17]. Here, by resolving these distributions across larval development, we additionally reveal how the strength of gravitaxis changes with age.

The heading distributions describe the dominant directions larval trajectories occupy, but not how fast the directions change. To quantify the timescale of this, we consider the trajectory-heading autocorrelation, 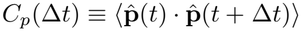 , where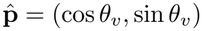, and fit

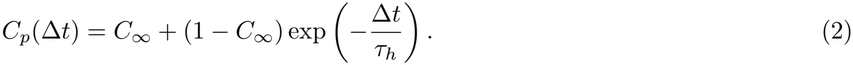

Here, *τ*_*h*_ characterizes relaxation toward the fitted plateau *C*_*∞*_, which measures the residual directional correlation (Fig. 3d). Our vertical-tracking observations reveal directional changes associated with helical turning and sustained oblique or horizontal bouts. We interpret *τ*_*h*_ alongside these observations: During 0–1 dpf, *C*_*p*_ remains close to unity throughout the 10-s fitting window, reflecting nearly monotonic buoyant rise. At approximately 2 dpf, *C*_*p*_ decreases rapidly toward a nonzero plateau, consistent with rapid heading fluctuations alongside a persistent directional bias. This rapid decorrelation accompanies the less regular swimming paths observed at this stage (SM Movie S1). At 3– 7 dpf, representative trajectories show organized helical bouts with a large spread of helix radius and pitch (Fig. 1e, 2d), indicating that short-time heading changes can also arise from repeated turning. At 8–9 dpf, the correlation plateau decreases and the relaxation becomes slower, with *τ*_*h*_ reaching its largest value among actively swimming stages at 9 dpf (Fig. 3d). Together with the broader heading distributions and sustained lateral excursions visible in the trajectories, these changes are consistent with longer-lived departures from vertical motion.

Together, these measurements summarize developmental changes in larval trajectories through three interpretable features: swimming speed, preferred direction, and directional persistence. Their consequences for vertical transport and horizontal spreading depend on their interaction with buoyancy and the surrounding flow.

### C. Developmental swimming in a fluctuating background flow

To examine these interactions, we construct a minimal model from the still-water measurements and expose each developmental stage to the same family of fluctuating flow fields. This allows us to test how developmental differences in swimming shape transport, and when flow-induced rotation overwhelms gravitactic alignment.

#### 1. A minimal model of gravitactic swimming

We first characterize larval transport over different timescales using the vertical and horizontal mean-squared displacements (MSDs; Fig. 4a). The vertical MSD remains approximately ballistic, MSD_*z*_ ∝ Δ*t*^2^, whereas the horizontal MSD approaches diffusive scaling at longer lags after active swimming begins, MSD_*x*_ ∝ Δ*t*, excluding the first two buoyant stages (0-1 dpf). This contrast is consistent with buoyancy and gravitactic swimming sustaining vertical transport while lateral velocity fluctuations decorrelate. To capture this gravitactic swimming, we construct a minimal model of a larva in the projected *x*–*z* plane with

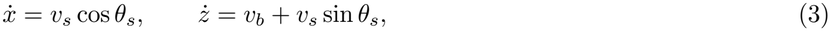

where (*x,z*) are the larva’s coordinates, *v*_*s*_ is the active swimming speed and *v*_*b*_ is the buoyant vertical drift. For actively swimming larvae, we infer the swimming heading as

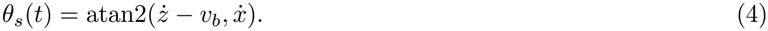

We interpret this buoyancy-corrected angle as a proxy for the direction of active propulsion. The Supplemental Material (SM, Sec. II.C) reports the corresponding *θ*_*s*_ distributions and compares the inferred headings with orientations independently extracted from the larval long axis in the videos (Figs. S2 and S3). We hold *v*_*s*_ and *v*_*b*_ constant within each trajectory. To focus on gravitaxis, we restrict the model to two active swimming modes, directed upward (*ψ*_+_ = +*π/*2) or downward (*ψ*_*−*_ = − *π/*2), with heading fluctuations around each preferred direction. In this minimal construction, we neglect switching between the two modes and assume that heading dynamics combine a restoring drift toward the preferred vertical direction with stochastic reorientation. In the aligned coordinate *η* = *θ*_*s*_ − *ψ*_*±*_, this gives (Fig. 4b),

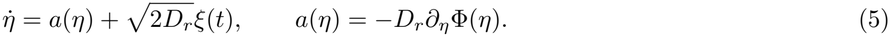

**FIG. 4.**
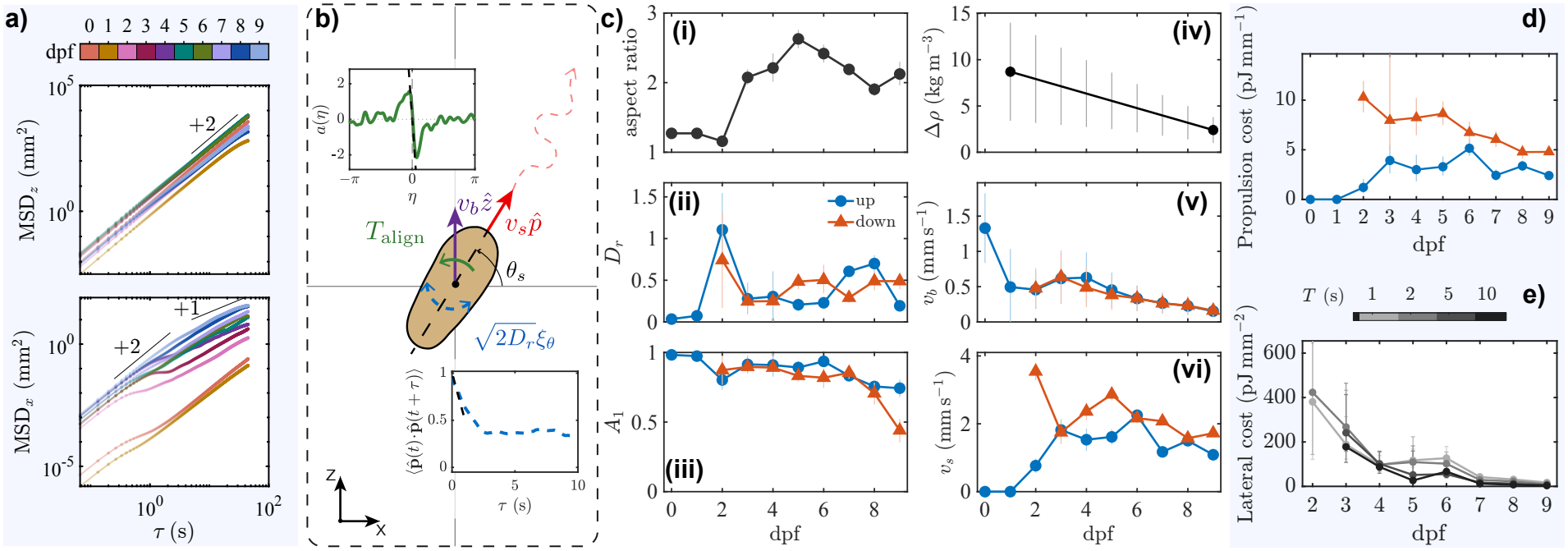
A minimal gravitactic model links orientation dynamics to dispersal. (a) Raw vertical and horizontal mean-squared displacements as functions of lag time. Persistent mean vertical motion produces an approximately ballistic vertical MSD, whereas finite heading memory produces approximately diffusive horizontal spreading after active swimming begins. (b) Minimal projected swimmer model with active speed *v*_*s*_, buoyant drift *v*_*b*_, and swimming heading *θ*_*s*_. Orientation is represented by a data-inferred restoring drift toward the preferred upward- or downward-directed state and stochastic reorientation with effective rotational diffusivity *D*_*r*_ . Insets show the fitted restoring drift and the short-lag swimming-heading autocorrelation *C*_*p,s*_ used to estimate *D*_*r*_ . (c) Stage-resolved model inputs: (i) body aspect ratio; (ii) *D*_*r*_ by swimming mode; (iii) first circular moment *A*_1_ of the swimming-heading distribution; (iv) buoyant density deficit Δ*ρ*, interpolated between the dpf 1 and dpf 9 passive-rise measurements; (v) buoyant drift *v*_*b*_; and (vi) active swimming speed *v*_*s*_ by mode. (d) Estimated propulsion cost per unit net vertical distance for straight upward (blue circles) and downward (orange triangles) swimming [Eq. (7)]. (e) Estimated propulsion cost per unit horizontal displacement variance [Eq. (8)], evaluated from experimental trajectories at lag times *T* = 1, 2, 5, and 10 s (gray scale). Estimates pool swimming modes with equal weight per contributing track. Error bars in panels (c, ii–vi), (d), and (e) show the ranges obtained by propagating the bounds on Δ*ρ*.

Here, *ξ*(*t*) is Gaussian white noise and *D*_*r*_ is an effective rotational diffusivity that sets the strength of stochastic heading fluctuations. We fit the orientation parameters separately for each developmental stage and swimming mode, assigning trajectories to the upward or downward mode according to the sign of their net vertical displacement, *z*(*t*_end_) − *z*(*t*_start_).

Within each mode, we assume stationary, Markovian heading dynamics with constant *D*_*r*_ and zero net probability current. The aligned headings define the conditional distribution *π*(*η* | *ψ* = *ψ*_*±*_), denoted by *π*(*η*) below. Under these assumptions, we infer the dimensionless alignment potential as Φ(*η*) = − log *π*(*η*) + const., giving the restoring drift *a*(*η*) = *D*_*r*_*∂*_*η*_ log *π*(*η*). Preferred headings therefore correspond to potential minima, toward which the drift directs the swimmer, without specifying a sensory or ciliary mechanism.

To determine how rapidly headings evolve within this potential, we calculate the swimming-heading autocorrelation, 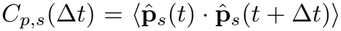, where 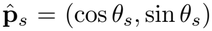 and the average includes only trajectories from the same developmental stage and swimming mode. Its initial decay gives (SM, Sec. III.A.1),

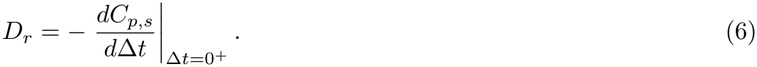

The heading distribution thus determines the shape of the alignment potential, while *D*_*r*_ sets the timescale of both restoring dynamics and stochastic reorientation. The model is mathematically analogous to Brownian motion in a potential well, with the balance between the restoring drift *a*(*η*) and rotational diffusion *D*_*r*_ determining the degree of gravitactic alignment. Here, the fitted drift and diffusion summarize the statistical heading changes associated with helical turning and departures from a regular helical path, but does not resolve the helical phase explicitly.

The parameters in Eqs. (3)–(5) were fit to experimental data conditioned on developmental stage (dpf) and swimming mode. At 0–1 dpf, we set *v*_*s*_ = 0, so the measured rise speed directly gives *v*_*b*_. For actively swimming stages, we estimate *v*_*b*_ from the measured larval geometry and density deficit, Δ*ρ* = *ρ*_fluid_ − *ρ*_body_, using the terminal force balance of an equivalent prolate spheroid [64]. We constrain Δ*ρ* using passive-rise measurements at 1 dpf and at 9 dpf after chemical arrest of ciliary propulsion, and linearly interpolate between these two stages (SM, Sec. II.B; Fig. S1). The direction of this interpolation is consistent with the developmental depletion of lipid reserves reported in other broadcast-spawning coral larvae [39, 40]. For each trajectory, we estimated *v*_*s*_ as the time-averaged magnitude of the velocity after subtracting *v*_*b*_ from its vertical component. Details of the fitting procedure and uncertainty propagation are given in the Supplemental Material, Sec. II.C.

Figure 4c summarizes the resulting developmental parameters. The aspect ratio increases as the initially rounded embryo develops into an elongated planula (Fig. 4c,i). The mode-resolved *D*_*r*_, which sets the short-time angular fluctuation rate, peaks near the onset of active swimming at 2 dpf (Fig. 4c,ii). To compare vertical alignment across stages, we use the first circular moment *A*_1_ = |⟨ *e*^*iη*^⟩ |= |⟨ cos⟩ *η*| for symmetric aligned distributions. Values near unity indicate headings concentrated around the preferred vertical direction, while a uniform distribution gives *A*_1_ = 0. Among actively swimming stages, *A*_1_ remains relatively high during 3–7 dpf and decreases at 8–9 dpf, indicating weaker vertical alignment without a systematic increase in *D*_*r*_ (Fig. 4c,ii–iii).

The density deficit decreases between the two measured developmental anchors [39, 40], and the inferred buoyant drift generally declines with age (Fig. 4c,iv–v). Active swimming speed varies with both development and swimming direction (Fig. 4c,vi). Differences between the upward- and downward-mode values of *v*_*s*_ remain after subtracting buoyancy, indicating that the strength of gravikinesis varies with age. Comparisons of the model with the measured heading distributions, autocorrelations, and MSDs are presented in SM, Sec. III.A.2 (Fig. S4).

Separating active propulsion from buoyant drift provides a basis for estimating the energetic cost of larval migration. Lipid reserves both support larval swimming and contribute to positive buoyancy [37, 38, 65]. Their depletion during development can reduce buoyant support, increasing reliance on active swimming to sustain upward transport [36, 39, 40]. Using the inferred swimming speeds, buoyant drifts, and body geometry, we estimate the propulsion cost per unit vertical distance (Fig. 4d). We approximate the propulsion power as 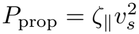, the power required to tow an equivalent prolate spheroid axially at speed *v*_*s*_. Over a time *t*, the estimated expenditure is *P*_prop_*t*. For straight vertical swimming, the net distance traveled is (*v*_*s*_ ± *v*_*b*_)*t*, giving

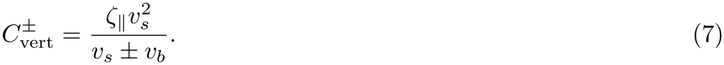

Here, *ζ*_*∥*_ is the axial drag coefficient, and the + and − signs denote upward and downward migration, respectively, using the fitted parameters for each swimming mode. We use this estimate to compare propulsion costs across development, rather than as a direct measure of ciliary or metabolic energy consumption.

At 0–1 dpf, buoyant ascent requires no active propulsion. The propulsion cost of upward migration rises after swimming begins and varies across the later stages. Downward migration has the higher estimated cost throughout the actively swimming stages, with buoyancy assisting ascent and opposing descent. Its cost nevertheless declines overall, reaching similar lowest values at 8–9 dpf. For a given active swimming speed and body geometry, weaker buoyancy allows faster net descent and therefore reduces the propulsion expenditure required to travel a fixed distance. This coupling between energy reserves, buoyancy, and swimming may help connect the changing demands of the coral life cycle. Positive buoyancy supports early dispersal without propulsion expenditure, while its decline as lipid reserves are consumed can make descent toward settlement habitats less costly later in development.

Even when swimming is predominantly gravity-aligned, oblique motion and horizontal bouts can contribute to lateral spreading. We therefore ask how much lateral spreading accompanies the observed swimming for a given estimated propulsion expenditure. We measure the horizontal displacement variance directly from the experimental trajectories over a common interval *T* , giving

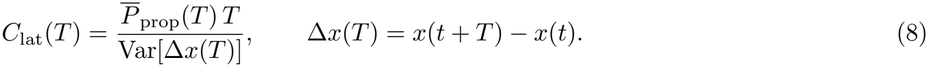

Here, 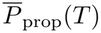 is the mean propulsion power of the trajectories contributing at each interval *T* , with equal weight per track. Lower values indicate more horizontal spreading for the same estimated propulsion expenditure. The lower costs at later developmental stages (Fig. 4e) therefore indicate that the increased lateral motion produces greater displacement variance relative to the estimated energy expended. Together with the broader heading distributions and longer directional persistence in Fig. 3, this suggests that development changes how effectively propulsion generates lateral spreading. Alongside the declining cost of descent, this shift may support the changing demands of exploration and settlement later in the larval stage.

#### 2. A fluctuating background flow progressively erodes gravitactic control

Our analysis so far has considered larval motion in still water. In nature however, coral larvae disperse through flows whose velocities can exceed their swimming speeds by orders of magnitude [46]. Gravitactic swimming can nevertheless bias their net vertical transport when realignment toward the preferred vertical direction is sufficiently rapid to counter flow-induced rotation [69]. More general navigation strategies can additionally show how swimmers can use environmental information to improve directed transport through complex or noisy flows [68, 70, 71]. Here, we test whether the developmental swimming states measured in still water continue to shape vertical transport and horizontal spreading, assuming that larvae respond only to gravity. Each fitted developmental stage is exposed to the same family of fluctuating velocity fields (Fig. 5a), retaining its measured swimming speed, buoyant drift, orientation dynamics, and body shape. The flow advects larvae and passively rotates them through local vorticity and strain (SM Movie S2).

**FIG. 5.**
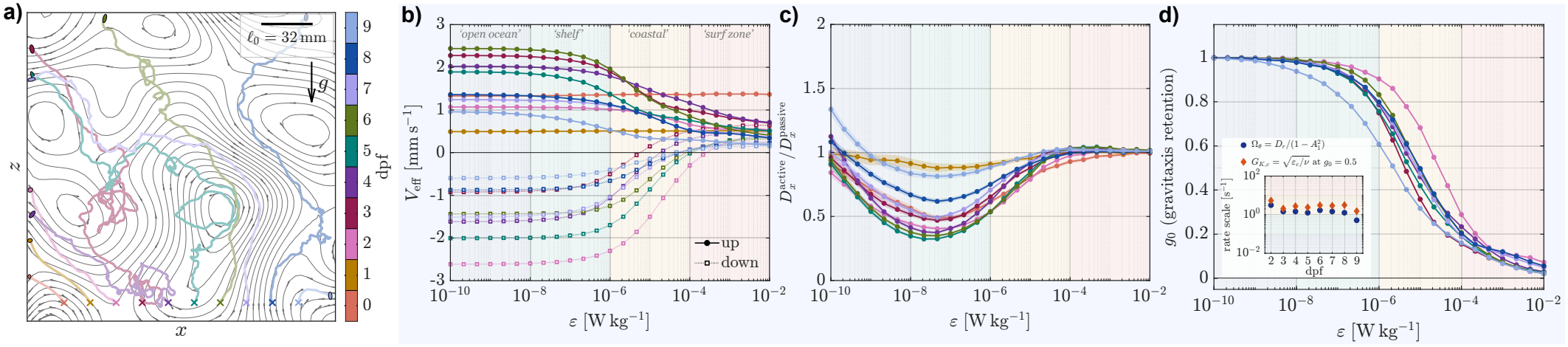
Developmental gravitactic control of dispersal in a fluctuating background flow. (a) Instantaneous stream-lines of the single-scale planar incompressible Gaussian random field at reference dissipation rate *ε* = 10^*−*8^ W kg^*−*1^, with one illustrative up-swimming trajectory per developmental stage. Crosses mark initial positions and enlarged ellipses mark endpoints; gravity points downward. Panels (b)–(d) use the time-evolving field. (b) Effective vertical velocity *V*_eff_ for each developmental stage and swimming mode. Solid and dotted curves denote upward- and downward-swimming modes. (c) Horizontal diffusivity relative to a passive tracer, 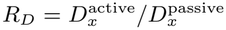, averaged equally over the two swimming modes. (d) Gravitaxis-retention index *g*_0_ = (*V*_eff_ −*v*_*b*_)*/*(*v*_*∞*_ − *v*_*b*_) for the actively swimming stages. The dotted line marks the half-retention crossover *g*_0_ = 0.5. Inset: still-water heading-relaxation rate 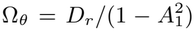 and the matched flow-gradient rate at this crossover. Background bands provide broad environmental context for the reference dissipation rate, although actual dissipation rates overlap among environments and vary with depth and time [66–68]. Shaded envelopes denote propagated standard errors across independent GRF realizations.

To connect these simulations to marine conditions, we use a reference three-dimensional turbulent dissipation rate *ε* as a single environmental parameter that sets the intensity and characteristic scales of a zero-mean, two-dimensional incompressible Gaussian random field in the observed *x*–*z* plane [72, 73]. For the fixed spatial and temporal closure used here, increasing *ε* increases the rms flow-speed scale as *u*_0_ ∝ *ε*^1*/*4^, while decreasing the spatial correlation length as *ℓ*_0_ ∝ *ε*^*−*1*/*4^ and the temporal correlation time as *τ*_0_ ∝ *ε*^*−*1*/*2^ (SM, Sec. III.B). The field is normalized so that its rms velocity-gradient magnitude matches the Kolmogorov gradient rate [67],

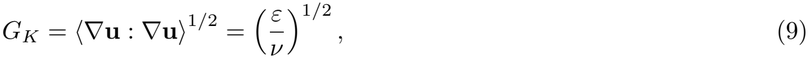

where *ν* is the kinematic viscosity and **u** = (*u*_*x*_, *u*_*z*_) is the background-flow velocity. Thus, larger *ε* represents a faster, finer-scale, and more rapidly decorrelating flow with stronger velocity gradients acting to reorient the larvae. This calibration allows an order-of-magnitude comparison with dissipation rates reported across marine environments, providing context for the environmental labels in (Fig. 5) [66–68]. Here, *ε* parameterizes a one-parameter family of statistically matched planar flows rather than denoting a dissipation rate generated dynamically by the planar field itself. The flow construction, numerical implementation, and sensitivity to the spatial and temporal closure scales are described in SM, Secs. III.B, III.D, and III.E, respectively.

In the flow, the larval dynamics become [74, 75]

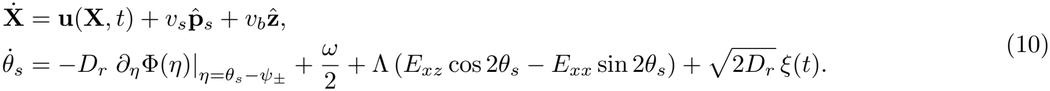

Where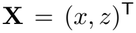, *ω* and *E*_*ij*_ are the planar vorticity and strain-rate components, and Λ = (*AR*^2^ − 1)*/*(*AR*^2^ + 1) is obtained from the measured aspect ratio. All biological parameters are fixed by Fig. 4; only the background-flow scales vary with *ε*.

To determine how developmental swimming influences transport in flow, we compare the effective vertical velocity, the horizontal diffusivity relative to a passive tracer, and a gravitaxis-retention index,

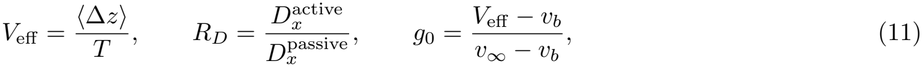

where *v*_*∞*_ = *v*_*b*_ + *v*_*s*_ ⟨ sin *θ*_*s*_ ⟩_0_ is the still-water vertical velocity. Here, *V*_eff_ measures mean ascent or descent over time *T* , while *R*_*D*_ *<* 1 and *R*_*D*_ *>* 1 indicate reduced and enhanced horizontal spreading, respectively, relative to neutrally buoyant passive tracers in the same flow. The index *g*_0_ normalizes vertical transport relative to the buoyant baseline by the still-water swimming contribution: *g*_0_ = 1 corresponds to the still-water mean vertical velocity, and *g*_0_ = 0 indicates transport at the buoyant baseline. This allows comparison across stages with different swimming speeds and buoyancies.

As expected, our numerical simulations indicate that buoyant ascent at 0–1 dpf remains nearly unchanged as *ε* increases (Fig. 5b). Thus, buoyancy supports upward transport without propulsion expenditure (Fig. 4d) and remains effective despite flow-induced reorientation. In contrast, the actively swimming stages approach their buoyant baselines as *ε* increases. Broadly speaking, intermediate stages ascend fastest at low *ε*, whereas the earliest active stage sustains net descent over the broadest range of *ε* (Fig. 5b). Swimming also modifies horizontal spreading within this model of upward- and downward-directed modes. At intermediate *ε, R*_*D*_ *<* 1 indicates reduced spreading relative to passive tracers, whereas *R*_*D*_ approaches unity at high *ε* (Fig. 5c). Increasing *ε* therefore shifts transport toward buoyant vertical drift and horizontal spreading comparable to that of passive tracers.

We next ask whether the effective gravitactic reorientation rate predicts the flow strength at which the swimming contribution is lost. We identify the half-retention crossover by 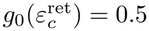 (Fig. 5d) and compare the corresponding flow-gradient rate with the effective still-water heading-relaxation rate (Fig. 5d, inset),

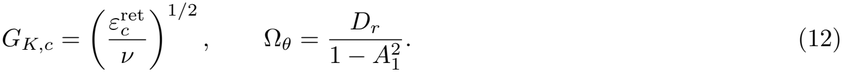

Here, Ω_*θ*_ is obtained by matching the initial slope and stationary plateau of the mode-resolved swimming-heading autocorrelation to the exponential form in Eq. (2), 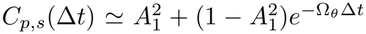. This gives 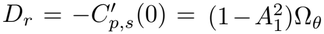 , with relaxation time 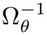 (SM, Sec. III.C). We note that this timescale describes *θ*_*s*_ within each swimming mode, whereas *τ*_*h*_ in Fig. 3d describes *θ*_*v*_ pooled across modes.

Across development, *G*_*K,c*_ has the same order of magnitude as Ω_*θ*_ (Fig. 5d, inset). Thus, the swimming contribution to vertical transport falls by half when the characteristic flow-gradient timescale, 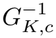 , becomes comparable to or shorter than the intrinsic heading-relaxation time,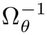. The corresponding gradient strengths are comparable to those reported in some coastal waters. Directed swimming retains a substantial contribution to vertical transport at lower *ε*, whereas stronger forcing at higher *ε* brings vertical transport toward the buoyant baseline and horizontal spreading toward the passive-tracer value. The environmental bands provide approximate context for this comparison, since dissipation rates vary with depth and time and overlap among environments [66, 67]. The precise crossover locations also depend on the spatial and temporal correlations of the imposed flow (SM, Sec. III.E; Fig. S5), and may shift when swimming is modeled in three dimensions or when larvae adjust their swimming in response to flow cues [68, 76]. Together, these results suggest that the measured still-water heading dynamics provide an estimate of the flow-gradient strength required to substantially reduce directed transport.

## IV. DISCUSSION

Our central finding is that *A. millepora* motility is a developmental program rather than a fixed trait. Over the first nine days post fertilization, larvae transition from passive, buoyancy-dominated rise, through a period of strongly gravity-aligned active swimming, and finally toward broader heading distributions and increased lateral motion (Figs. 2 and 3). These later changes may support the search for suitable settlement sites, a behavior documented in competent *A. millepora* larvae [77]. Our minimal model integrates the accompanying changes in morphology, buoyancy, active propulsion, gravitactic alignment, and reorientation to describe how their changing balance shapes larval transport across development (Fig. 4).

The propulsion-cost estimates suggest how this developmental progression may relate to the finite energy reserves available before settlement [78] (Fig. 4d,e). Lipid reserves support early buoyant ascent without propulsion expenditure, but that same buoyancy opposes descent once active swimming develops. Declining buoyant support contributes to the overall reduction in the estimated cost of straight downward migration. Later-stage trajectories also generate greater horizontal displacement variance per unit estimated propulsion expenditure. Together, these trends suggest that descent and lateral movement may become less costly as larvae draw on their reserves. These comparisons use an equivalent-body estimate of propulsion power; direct measurements would be needed to establish the corresponding ciliary and metabolic costs.

Coral larvae have previously been described as poor swimmers because their swimming speeds are substantially lower than characteristic water velocities measured on and around reefs [46]. Our simulations assess their baseline capacity to maintain a vertical swimming bias while being carried and reoriented by flow. Our model captures the observed gravitactic alignment without requiring knowledge of its sensory or mechanical origin. Passive reorientation arising from asymmetries in body shape or internal density has been observed, but its contribution relative to active regulation of coral larval orientation remains unresolved [17]. Similarly, turbulence experiments have not resolved whether reduced vertical swimming reflects disrupted orientation, altered swimming activity, or both [44]. We assume that the intrinsic orientation dynamics measured in still water remain unchanged, with no additional behavioral response to flow cues. Under these assumptions, swimming speed relative to current speed is not the only relevant comparison. Comparing the effective heading-relaxation rate Ω_*θ*_ with the imposed gradient rate, such as *G*_*K*_ ∼ (*ε/ν*)^1*/*2^ used here, provides an approximate scale for when flow-induced rotation can disrupt the vertical bias (Fig. 5). Under the approximate environmental mapping used here, the simulations predict substantial retention of directed swimming under weaker open-ocean- and shelf-like gradients, with progressively reduced swimming contributions under stronger coastal- and surf-zone-like gradients.

Maintaining a vertical bias could therefore influence dispersal even when swimming speeds are small relative to ambient currents. Persistent changes in depth can move larvae between flow layers, altering their horizontal advection [79], residence time, and exposure to the benthic boundary layer [49–51]. Our results suggest that the capacity to produce these changes varies across development. Early buoyancy provides an upward bias that persists despite flow-induced reorientation, while active swimming subsequently permits descent and strengthens upward transport during intermediate stages under weak flow. Later weakening of gravitactic alignment and increased lateral motion may support the search for settlement sites. Near the benthos, flow influences larval delivery and successful attachment [80–82], while reef topography can further promote substrate encounter by reducing local flow speeds and redirecting larvae toward settlement surfaces [46, 47, 54]. Field experiments also show that settlement-surface design and microbial coatings can enhance coral larval settlement [83]. Developmental swimming and local hydrodynamic structure may thus act together to influence arrival at suitable habitats, although these ecological roles remain hypotheses.

Our two-dimensional, zero-mean GRF provides a controlled comparison of how developmental swimming states respond to increasing flow gradients. It omits mean currents, depth-dependent flow structure, waves, and substrate interactions, and therefore cannot directly predict reef connectivity or settlement. The reference dissipation rate *ε* matches the rms planar velocity-gradient magnitude to a three-dimensional Kolmogorov scale, without reproducing the turbulent cascade or intermittency. Incorporating the measured developmental behaviors into more realistic reef flows, including models that resolve wave-driven flow over benthic topography [47, 54], would help connect these results to substrate encounter and settlement. Reports of altered swimming following mechanical disturbance motivate testing how behavioral responses to flow modify these predictions [35]. Ciliary sensory roles proposed for *Nematostella* larvae and polycystin-1-dependent mechanosensation demonstrated in its polyps provide candidate mechanisms for investigation, although their involvement in coral larval flow-gradient detection remains unresolved [84, 85].

Our results show how development changes the balance between buoyancy and active swimming, shaping how larvae interact with their physical environment across developmental stages. Successive larval stages may serve different ecological functions, with early buoyant ascent supporting dispersal and later descent and lateral movement facilitating the search for settlement habitats. Natural selection may therefore shape the timing and sequence of these behaviors, alongside morphology, to meet changing demands across development. This perspective places developmental changes in behavior at the center of efforts to connect individual larval motion to ecological outcomes. Incorporating these changes and their interaction with flow into dispersal models could improve predictions of transport between reefs and encounters with suitable settlement habitat, informing restoration strategies that depend on larval connectivity and successful recruitment.

## V. ACKNOWLEDGMENTS

The authors thank Rebecca Konte for the illustrations used in Fig. 1. Computations for this project were performed in part on the Sherlock cluster. We thank Stanford University and the Stanford Research Computing Center for providing computational resources and support. Part of this work was performed at the Cell Sciences Imaging Facility (RRID: SCR 017787) at Stanford University, and we thank the assistance provided by D. Lenzi. Q.Z. and M.P. acknowledge support from Human Frontier Science Program and National Science Foundation (NSF) (grant no. 135316). M.P. acknowledges further financial support from the Schmidt Futures Innovation Fellowship, the Moore Foundation, the Dalio Foundation, the NSF Center for Cellular Construction (grant no. DBI-1548297), ARIA Nature Computes Better Program, and the Woods Institute for the Environment.

## VI. AUTHOR CONTRIBUTIONS

M.P., I.H., Q.Z., and B.F. designed the study. E.L.N., S.M. and R.A. carried out coral spawning. Q.Z. and I.H. performed a majority of the tracking microscopy experiments, and P.V. performed the immunostaining. I.H. analyzed the data and constructed the minimal model, with feedback from B.F and M.P. I.H. and B.F. wrote the manuscript, with feedback from all authors.

## VII. DATA AVAILABILITY

The raw trajectory data, videos, and code are available from the corresponding author upon request.

## Notes

### Competing Interest Statement

The authors have declared no competing interest.

## References

[1] R. K. Cowen, G. Gawarkiewicz, J. Pineda, S. R. Thorrold, and F. E. Werner, Population connectivity in marine systems an overview, Oceanography 20, 14 (2007).

[2] J. Veron, M. Stafford-Smith, L. DeVantier, and E. Turak, Overview of distribution patterns of zooxanthellate Scleractinia, Frontiers in Marine Science 1, 10.3389/fmars.2014.00081 (2015).

[3] S. E. Kahng, J. R. Garcia-Sais, H. L. Spalding, E. Brokovich, D. Wagner, E. Weil, L. Hinderstein, and R. J. Toonen, Community ecology of mesophotic coral reef ecosystems, Coral Reefs 29, 255 (2010).

[4] J. M. Roberts, A. J. Wheeler, and A. Freiwald, Reefs of the Deep: The Biology and Geology of Cold-Water Coral Ecosystems, Science 312, 543 (2006).

[5] S. D. Cairns, Deep-water corals: an overview with special reference to diversity and distribution of deep-water scleractinian corals, Bulletin of marine Science 81, 311 (2007).

[6] J.-C. Ortiz, N. H. Wolff, K. R. N. Anthony, M. Devlin, S. Lewis, and P. J. Mumby, Impaired recovery of the Great Barrier Reef under cumulative stress, Science Advances 4, eaar6127 (2018).

[7] J. Morais, S. B. Tebbett, R. A. Morais, and D. R. Bellwood, Natural recovery of corals after severe disturbance, Ecology Letters 27, e14332 (2024).

[8] S. M. Rosales, B. D. Young, A. J. Bright, E. Montes, J.-Z. Zhang, N. Traylor-Knowles, and D. E. Williams, Reef site and habitat influence effectiveness of Acropora palmata restoration and its microbiome in the Florida Keys, Communications Earth & Environment 5, 689 (2024).

[9] L. Chapron, I. B. Kuffner, D. W. Kemp, A. M. Hulver, E. F. Keister, A. Stathakopoulos, L. A. Bartlett, E. O. Lyons, and A. G. Grottoli, Heterotrophy, microbiome, and location effects on restoration efficacy of the threatened coral Acropora palmata, Communications Earth & Environment 4, 233 (2023).

[10] C. Doropoulos, G. Roff, G. Carlin, M. Gouezo, D. Dela Cruz, A. Chai, L. Hardiman, L. Hasson, D. P. Thomson, and P. L. Harrison, Larval seedboxes: A modular and effective tool for scaling coral reef restoration, Ecological applications 35, e70140 (2025).

[11] R. K. Cowen, K. M. Lwiza, S. Sponaugle, C. B. Paris, and D. B. Olson, Connectivity of marine populations: open or closed?, Science 287, 857 (2000).

[12] S. Sponaugle, R. K. Cowen, A. Shanks, S. G. Morgan, J. M. Leis, J. Pineda, G. W. Boehlert, M. J. Kingsford, K. C. Lindeman, C. Grimes, et al., Predicting self-recruitment in marine populations: biophysical correlates and mechanisms, Bulletin of Marine Science 70, 341 (2002).

[13] G. F. Ricardo, R. J. Jones, A. P. Negri, and R. Stocker, That sinking feeling: Suspended sediments can prevent the ascent of coral egg bundles, Scientific Reports 6, 21567 (2016).

[14] G. F. Ricardo, R. J. Jones, M. Nordborg, and A. P. Negri, Settlement patterns of the coral acropora millepora on sediment-laden surfaces, Science of the Total Environment 609, 277 (2017).

[15] M. Ishikawa and M. Hatta, Identification of pre-settling behavior leading the planula larva to settlement in the coral acropora, Zoological Science 43 (2026).

[16] A. M. Szmant and M. G. Meadows, Developmental changes in coral larval buoyancy and vertical swimming behavior: implications for dispersal and connectivity, in Proc 10th Int Coral Reef Symp, Vol. 1 (2006) pp. 431–437.

[17] A. Takeda-Sakazume, J. Honjo, S. Sasano, K. Matsushima, S. A. Baba, Y. Mogami, and M. Hatta, Gravitactic Swimming of the Planula Larva of the Coral Acropora: Characterization of Straightforward Vertical Swimming, Zoological Science 40, 44 (2022).

[18] A. I. Larsson, J. Jarnegren, S. M. Strömberg, M. P. Dahl, T. Lundälv, and S. Brooke, Embryogenesis and larval biology of the cold-water coral lophelia pertusa, PLoS One 9, e102222 (2014).

[19] S. Weinberg, The light-dependent behaviour of planula larvae of eunicella singularis and corallium rubrum and its implication for octocorallian ecology, Bijdragen tot de Dierkunde 49, 16 (1979).

[20] M. Lønnum, J. Hovland, M. M. Schuldt, E. S. Nilssen, J. Davila-Velderrain, J. Järnegren, and L. van Giesen, Dispersal behavior in a cold-water coral is orchestrated via stage and species-specific physiology, bioRxiv , 2026 (2026).

[21] R. N. Poon, T. A. Westwood, H. Laeverenz-Schlogelhofer, E. Brodrick, J. Craggs, E. E. Keaveny, G. Jékely, and K. Y. Wan, Ciliary propulsion and metachronal coordination in reef coral larvae, Physical Review Research 5, L042037 (2023).

[22] B. D. Limer, O. Payne, A. Brancato, M. Mitchell, C. Abu-Kubie, J. Figueiredo, and D. M. Holstein, Life history and early ontogeny determine vertical swimming behaviors in the larvae of caribbean corals, Journal of Experimental Marine Biology and Ecology 578, 152035 (2024).

[23] S. Brooke and C. Young, Embryogenesis and larval biology of the ahermatypic scleractinian oculina varicosa, Marine Biology 146, 665 (2005).

[24] M. Vermeij, N. D. Fogarty, and M. Miller, Pelagic conditions affect larval behavior, survival, and settlement patterns in the caribbean coral montastraea faveolata, Marine Ecology Progress Series 310, 119 (2006).

[25] S. M. Strömberg and A. I. Larsson, Larval behavior and longevity in the cold-water coral lophelia pertusa indicate potential for long distance dispersal, Frontiers in Marine Science 4, 411 (2017).

[26] M. J. Kingsford, J. M. Leis, A. Shanks, K. C. Lindeman, S. G. Morgan, and J. Pineda, SENSORY ENVIRONMENTS, LARVAL ABILITIES AND LOCAL SELF-RECRUITMENT, BULLETIN OF MARINE SCIENCE 70 (2002).

[27] M. J. A. Vermeij, K. L. Marhaver, C. M. Huijbers, I. Nagelkerken, and S. D. Simpson, Coral Larvae Move toward Reef Sounds, PLOS ONE 5, e10660 (2010).

[28] P. T. Raimondi and A. N. Morse, The consequences of complex larval behavior in a coral, Ecology 81, 3193 (2000).

[29] L. Harrington, K. Fabricius, G. De’Ath, and A. Negri, Recognition and selection of settlement substrata determine post-settlement survival in corals, Ecology 85, 3428 (2004).

[30] A. J. Mulla, C.-H. Lin, S. Takahashi, and Y. Nozawa, Photo-movement of coral larvae influences vertical positioning in the ocean, Coral Reefs 40, 1297 (2021).

[31] M. E. Strader, S. W. Davies, and M. V. Matz, Differential responses of coral larvae to the colour of ambient light guide them to suitable settlement microhabitat, Royal Society open science 2, 150358 (2015).

[32] Y. Sakai, K. Kato, H. Koyama, A. Kuba, H. Takahashi, T. Fujimori, M. Hatta, A. P. Negri, A. H. Baird, and N. Ueno, A step-down photophobic response in coral larvae: implications for the light-dependent distribution of the common reef coral, acropora tenuis, Scientific Reports 10, 17680 (2020).

[33] D. F. Gleason and D. K. Hofmann, Coral larvae: from gametes to recruits, Journal of Experimental Marine Biology and Ecology 408, 42 (2011).

[34] J. W. Pysanczyn, E. A. Williams, E. Brodrick, D. Robert, J. Craggs, K. L. Marhaver, and S. D. Simpson, The role of acoustics within the sensory landscape of coral larval settlement, Frontiers in Marine Science 10, 1111599 (2023).

[35] K. Guizien, N. Viladrich, Á. Martínez-Quintana, and L. Bramanti, Survive or swim: different relationships between migration potential and larval size in three sympatric mediterranean octocorals, Scientific reports 10, 18096 (2020).

[36] E. M. Graham, A. H. Baird, S. R. Connolly, M. A. Sewell, and B. L. Willis, Rapid declines in metabolism explain extended coral larval longevity, Coral Reefs 32, 539 (2013).

[37] J. L. Matthews, N. Bartels, H. Kish, L. La Motta, M. P. Padula, D. J. Hughes, Z. J. Crane, C. Humphrey, N. Boulotte, C. Roper, M. J. H. van Oppen, J.-B. Raina, and D. J. Suggett, Sterols are key to coral larvae survival, swimming capacity, and thermal tolerance, Communications Biology 8, 1494 (2025).

[38] I. Arai, M. Kato, A. Heyward, Y. Ikeda, T. Iizuka, and T. Maruyama, Lipid composition of positively buoyant eggs of reef building corals, Coral Reefs 12, 71 (1993).

[39] J. Figueiredo, A. H. Baird, M. F. Cohen, J.-F. Flot, T. Kamiki, T. Meziane, M. Tsuchiya, and H. Yamasaki, Ontogenetic change in the lipid and fatty acid composition of scleractinian coral larvae, Coral Reefs 31, 613 (2012).

[40] S. Harii, K. Nadaoka, M. Yamamoto, and K. Iwao, Temporal changes in settlement, lipid content and lipid composition of larvae of the spawning hermatypic coral Acropora tenuis, Marine Ecology Progress Series 346, 89 (2007).

[41] C. J. Randall, C. Giuliano, B. Stephenson, T. N. Whitman, C. A. Page, E. A. Treml, M. Logan, and A. P. Negri, Larval precompetency and settlement behaviour in 25 Indo-Pacific coral species, Communications Biology 7, 1 (2024).

[42] S. R. Connolly and A. H. Baird, Estimating dispersal potential for marine larvae: Dynamic models applied to scleractinian corals, Ecology 91, 3572 (2010).

[43] R. M. Attenborough, D. C. Hayward, U. Wiedemann, S. Forêt, D. J. Miller, and E. E. Ball, Expression of the neuropeptides RFamide and LWamide during development of the coral Acropora millepora in relation to settlement and metamorphosis, Developmental Biology 446, 56 (2019).

[44] V. Fagerström, G. Brostrom, and A. I. Larsson, Turbulence affects larval vertical swimming in the cold-water coral lophelia pertusa, Frontiers in Marine Science 9, 1062884 (2022).

[45] E. Wolanski and W. M. Hamner, Topographically controlled fronts in the ocean and their biological influence, Science 241, 177 (1988).

[46] T. Hata, J. S. Madin, V. R. Cumbo, M. Denny, J. Figueiredo, S. Harii, C. J. Thomas, and A. H. Baird, Coral larvae are poor swimmers and require fine-scale reef structure to settle, Scientific Reports 7, 2249 (2017).

[47] D. Gysbers, M. A. Levenstein, and G. Juarez, Effect of substrate topography on benthic boundary layer flow: Implications for marine larval transport and settlement, Physical Review Fluids 10, 064501 (2025).

[48] B. L. Willis and J. K. Oliver, Direct tracking of coral larvae: implications for dispersal studies of planktonic larvae in topographically complex environments, Ophelia 32, 145 (1990).

[49] H. L. Fuchs, E. J. Hunter, E. L. Schmitt, and R. A. Guazzo, Active downward propulsion by oyster larvae in turbulence, Journal of Experimental Biology , jeb.079855 (2012).

[50] H. L. Fuchs, G. P. Gerbi, E. J. Hunter, and A. J. Christman, Waves cue distinct behaviors and differentiate transport of congeneric snail larvae from sheltered versus wavy habitats, Proceedings of the National Academy of Sciences 115, E7532 (2018).

[51] J. D. Wheeler, E. Secchi, R. Rusconi, and R. Stocker, Not Just Going with the Flow: The Effects of Fluid Flow on Bacteria and Plankton, Annual Review of Cell and Developmental Biology 35, 213 (2019).

[52] C. B. Paris and R. K. Cowen, Direct evidence of a biophysical retention mechanism for coral reef fish larvae, Limnology and Oceanography 49, 1964 (2004).

[53] J. Greenwood, C. Sun, C. Doropoulos, D. Thomson, M. Baird, J. Porobic, and S. Condie, Passive retention of simulated larvae on coral reefs, Royal Society open science 12, 241708 (2025).

[54] M. A. Levenstein, D. J. Gysbers, K. L. Marhaver, S. Kattom, L. Tichy, Z. Quinlan, H. M. Tholen, L. Wegley Kelly, M. J. Vermeij, A. J. Wagoner Johnson, et al., Millimeter-scale topography facilitates coral larval settlement in wave-driven oscillatory flow, PLoS One 17, e0274088 (2022).

[55] A. Metaxas and M. Saunders, Quantifying the “bio-” components in biophysical models of larval transport in marine benthic invertebrates: advances and pitfalls, The Biological Bulletin 216, 257 (2009).

[56] Y. C. Tay, J. R. Guest, L. M. Chou, and P. A. Todd, Vertical distribution and settlement competencies in broadcast spawning coral larvae: Implications for dispersal models, Journal of Experimental Marine Biology and Ecology 409, 324 (2011).

[57] W. Thielicke and R. Sonntag, Particle image velocimetry for matlab: Accuracy and enhanced algorithms in pivlab, Journal of Open Research Software 9, 12 (2021).

[58] D. Yuan, N. Nakanishi, D. K. Jacobs, and V. Hartenstein, Embryonic development and metamorphosis of the scyphozoan aurelia, Development genes and evolution 218, 525 (2008).

[59] D. Krishnamurthy, H. Li, F. Benoit du Rey, P. Cambournac, A. G. Larson, E. Li, and M. Prakash, Scale-free vertical tracking microscopy, Nature Methods 17, 1040 (2020).

[60] E. A. Brodrick, K. Jokura, J. Craggs, R. Poon, H. Laeverenz-Schlogelhofer, K. Y. Wan, and G. Jékely, A light-off response characterised by body contraction and ciliary arrest in acropora coral larvae, Journal of Experimental Biology 229, jeb252360 (2026).

[61] B. Li, J. Yan, W. Wu, Z. Zhu, and X. Hu, High Performance Visual Tracking with Siamese Region Proposal Network, in 2018 IEEE/CVF Conference on Computer Vision and Pattern Recognition (IEEE, Salt Lake City, UT, 2018) pp. 8971–8980.

[62] G. F. Ricardo, R. J. Jones, P. L. Clode, and A. P. Negri, Mucous secretion and cilia beating defend developing coral larvae from suspended sediments, PLoS One 11, e0162743 (2016).

[63] D. C. Hayward, J. Catmull, J. S. Reece-Hoyes, H. Berghammer, H. Dodd, S. J. Hann, D. J. Miller, and E. E. Ball, Gene structure and larval expression of cnox-2am from the coral acropora millepora, Development genes and evolution 211, 10 (2001).

[64] A. T. Chwang and T. Y.-T. Wu, Hydromechanics of low-reynolds-number flow. part 2. singularity method for stokes flows, Journal of Fluid mechanics 67, 787 (1975).

[65] S. Harii, M. Yamamoto, and O. Hoegh-Guldberg, The relative contribution of dinoflagellate photosynthesis and stored lipids to the survivorship of symbiotic larvae of the reef-building corals, Marine Biology 157, 1215 (2010).

[66] H. L. Fuchs and G. P. Gerbi, Seascape-level variation in turbulence-and wave-generated hydrodynamic signals experienced by plankton, Progress in Oceanography 141, 109 (2016).

[67] P. J. Franks, B. G. Inman, J. A. MacKinnon, M. H. Alford, and A. F. Waterhouse, Oceanic turbulence from a planktonic perspective, Limnology and Oceanography 67, 348 (2022).

[68] R. Monthiller, A. Loisy, M. A. Koehl, B. Favier, and C. Eloy, Surfing on turbulence: a strategy for planktonic navigation, Physical Review Letters 129, 064502 (2022).

[69] W. M. Durham, J. O. Kessler, and R. Stocker, Disruption of vertical motility by shear triggers formation of thin phytoplankton layers, science 323, 1067 (2009).

[70] L. Piro, B. Mahault, and R. Golestanian, Optimal navigation of microswimmers in complex and noisy environments, New Journal of Physics 24, 093037 (2022).

[71] L. Piro, A. Vilfan, R. Golestanian, and B. Mahault, Energetic cost of microswimmer navigation: The role of body shape, Physical Review Research 6, 013274 (2024).

[72] J. Bec, K. Gustavsson, and B. Mehlig, Statistical models for the dynamics of heavy particles in turbulence, Annual Review of Fluid Mechanics 56, 189 (2024).

[73] K. Gustavsson, F. Berglund, P. Jonsson, and B. Mehlig, Preferential sampling and small-scale clustering of gyrotactic microswimmers in turbulence, Physical review letters 116, 108104 (2016).

[74] T. Pedley and J. O. Kessler, Hydrodynamic phenomena in suspensions of swimming microorganisms, Annual Review of Fluid Mechanics 24, 313 (1992).

[75] G. B. Jeffery, The motion of ellipsoidal particles immersed in a viscous fluid, Proceedings of the Royal Society of London. Series A, Containing papers of a mathematical and physical character 102, 161 (1922).

[76] M. H. DiBenedetto, R. Monthiller, C. Eloy, and L. S. Mullineaux, Plankton active response to turbulence enables efficient transport, Journal of Experimental Biology 228, jeb251123 (2025).

[77] F. J. Pollock, S. M. Katz, J. A. van de Water, S. W. Davies, M. Hein, G. Torda, M. V. Matz, V. H. Beltran, P. Buerger,E. Puill-Stephan, et al., Coral larvae for restoration and research: a large-scale method for rearing acropora millepora larvae, inducing settlement, and establishing symbiosis, PeerJ 5, e3732 (2017).

[78] D. E. Wendt, Energetics of larval swimming and metamorphosis in four species of bugula (bryozoa), The Biological Bulletin 198, 346 (2000).

[79] S. F. Gary, A. D. Fox, A. Biastoch, J. M. Roberts, and S. A. Cunningham, Larval behaviour, dispersal and population connectivity in the deep sea, Scientific reports 10, 10675 (2020).

[80] C. Butman, Larval settlement of soft-sediment invertebrates: the spatial scales of pattern explained by active habitat selection and the emerging role of hydrodynamical processes, Oceanogr Mar Biol Annu Rev 25, 113 (1987).

[81] A. Abelson and M. Denny, Settlement of marine organisms in flow, Annual Review of Ecology and Systematics 28, 317 (1997).

[82] M. A. Reidenbach, J. R. Koseff, and M. Koehl, Hydrodynamic forces on larvae affect their settlement on coral reefs in turbulent, wave-driven flow, Limnology and oceanography 54, 318 (2009).

[83] A. Thode, N. Levy, O. Boulais, S. Kundu, M. Freckelton, D. Schar, M. Hadfield, J. Reichert, H. Jorissen, N. Kramer, et al., In situ enhancement of coral larval settlement using acoustic enrichment, engineered habitats and living materials, Communications Biology 9, 1288 (2026).

[84] M. Lønnum, M. M. Schuldt, J. Hovland, J. Davila-Velderrain, and L. Van Giesen, Developmentally timed sensory integration enables efficient larval dispersal, Iscience 29 (2026).

[85] J. Baranyk, K. Malir, M. A. Silva, S. Rieck, G. Scheve, and N. Nakanishi, Structural, molecular and developmental evidence for cell-type diversity in cnidarian mechanosensory neurons, Nature Communications 16, 1514 (2025).

