## Supplemental Material for "Behavioral Plasticity and Dispersal of Free Swimming Coral Larvae via Vertical Tracking Microscopy"

[1]*Department of Bioengineering, Stanford University, Stanford CA 94305 USA*
[2]*Institute for Biodiversity Science and Sustainability, California Academy of Sciences, San Francisco, California*
[3]*Daniel P. Haerther Center for Conservation and Research, John G. Shedd Aquarium, Chicago, IL, USA*
(Dated: November 19, 2026)

### CONTENTS

I. Supplemental Movies 1

II. Experimental Methods 2
A. Data summary 2
B. Inference of larval density from buoyant rise 3
C. Separation of buoyant drift and active swimming speed 4
D. Fixation and immunostaining 5

III. Theoretical model 6
A. Minimal projected swimmer model 6
1. Rotational diffusivity from the heading autocorrelation 6
2. Numerical validation of the minimal model 7
B. Spectral construction and normalization of the Gaussian random flow 7
C. Effective heading-relaxation rate 8
D. Numerical implementation of transport simulations 9
E. Sensitivity to the GRF closure parameters 10

References 11

### I. SUPPLEMENTAL MOVIES

**Supplemental Movie S1. Developmental changes in larval motion.** Representative vertical-tracking recordings of individual *Acropora millepora* larvae at 0, 1, 2, 4, 5, 7, 8, and 9 dpf. Each sequence pairs the microscopic view (left) with the corresponding trajectory (right), colored by speed; the yellow marker indicates the position corresponding to the displayed time. The sequences illustrate buoyant rise at 0–1 dpf, the onset of active swimming at 2 dpf, organized gravitactic swimming and helical trajectories at intermediate stages, and oblique and late stage swimming at 8–9 dpf. Elapsed experimental time is shown above each microscopic view. Trajectory coordinates are in millimeters and speeds in $\mathrm{mm\,s^{-1}}$.

**Supplemental Movie S2. Gravitactic swimming in fluctuating background flow.** Illustrative simulations of the fitted 7-dpf upward-swimming mode at reference dissipation rates $\varepsilon = 10^{-8}$, $10^{-6}$, and $10^{-4}\,\mathrm{W\,kg^{-1}}$ (left to right). Swimmers are advected and passively rotated by a two-dimensional Gaussian random flow while retaining their fitted buoyant drift, propulsion speed, and intrinsic heading dynamics. No flow sensing or switching between swimming modes is included. The upper panels show fixed laboratory-frame windows with the same physical dimensions. Ellipses indicate swimming headings and retain the fitted aspect ratio, with size enlarged for visibility. Gray curves and arrows indicate instantaneous flow streamlines and directions. Display density is maintained by replacing swimmers that leave the viewing window. The middle row shows the instantaneous swimming contribution to mean vertical velocity, $v_s\langle\sin\theta_s\rangle$ (blue), and the bottom row shows the instantaneous total mean vertical velocity, $\langle\dot{z}\rangle = \langle u_z\rangle + v_b + v_s\langle\sin\theta_s\rangle$ (orange). Dashed lines of matching colors show the corresponding still-water expectations. The total velocity includes buoyant drift and the vertical fluid velocity sampled along the trajectories.

---

* These authors contributed equally.

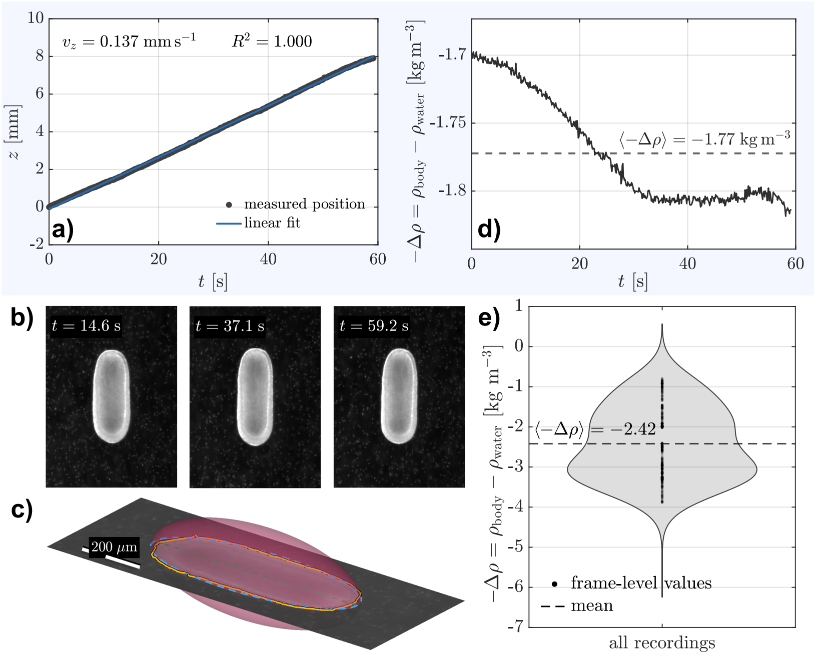


FIG. S1. **Inferring larval density from passive vertical motion and a prolate-spheroid body model.** (a) Vertical position $z$ of a representative larva during a buoyancy measurement immediately after $MgCl_2$ treatment. The larva rose nearly linearly over the 60-s observation interval; a linear fit gave a vertical velocity $v_z = 0.137$ mm s$^{-1}$ with $R^2 = 1.000$. (b) Corresponding images of the larva at $t = 14.6$, 37.1, and 59.2 s, showing that its body shape and predominantly vertical orientation remained approximately constant during the rise. (c) Prolate-spheroid representation used in the density inference for a representative image acquired at $t = 29.8$ s. The orange contour shows the segmented larval silhouette, the blue dashed curve shows the equivalent in-plane ellipse defined by the measured major and minor axes, and the translucent surface shows the corresponding prolate spheroid. The green dashed line indicates the major axis and the assumed direction of axial translation. Scale bar, 200 $\mu$m. (d) Frame-resolved density contrast, $\rho_{\mathrm{body}} - \rho_{\mathrm{water}} = -\Delta\rho$, obtained by combining the measured rise velocity with the corresponding prolate-spheroid geometry and axial hydrodynamic resistance. The dashed line denotes the mean for this recording. (e) Distribution of frame-level density estimates across all analyzed larval recordings. The violin width represents the estimated probability density, black points show frame-level measurements, and the dashed line denotes the mean of the included measurements.

### II. EXPERIMENTAL METHODS

#### A. Data summary

TABLE S1. Trajectory sampling across larval development. $N$ denotes the number of trajectories recorded at each nominal day post fertilization (dpf). Measurement windows indicate the range of acquisition times within each developmental day.

| dpf | Date | $N$ | Measurement window | Window span (h) |
|---|---|---|---|---|
| 0 | 2022-12-18 | 21 | 12:46:12–00:05:57 | 11.33 |
| 1 | 2022-12-19 | 8 | 10:58:05–23:52:45 | 12.91 |
| 2 | 2022-12-20 | 11 | 15:38:05–22:28:34 | 6.84 |
| 3 | 2022-12-21 | 20 | 19:10:45–23:44:06 | 4.56 |
| 4 | 2022-12-22 | 29 | 23:33:56–11:22:48 | 11.81 |
| 5 | 2022-12-23 | 55 | 12:00:32–00:35:34 | 12.58 |
| 6 | 2022-12-24 | 31 | 12:28:42–23:40:58 | 11.20 |
| 7 | 2022-12-25 | 97 | 12:22:37–21:02:41 | 8.67 |
| 8 | 2022-12-26 | 41 | 09:22:56–00:46:20 | 15.39 |
| 9 | 2022-12-27 | 32 | 17:01:15–19:31:41 | 2.51 |

#### B. Inference of larval density from buoyant rise

Larval buoyancy was measured at two developmental anchor points, dpf 1 and dpf 9, using a similar analysis as Takeda-Sakazume et al. [1] but different experimental conditions to isolate passive vertical motion. At 1 dpf, weak ciliary beating was visible; however, the larvae followed nearly monotonic, predominantly vertical trajectories, with no obvious change in path or rise speed attributable to active propulsion. We therefore treated the measured terminal rise speed as the buoyant rise speed of the larva. At 9 dpf, larvae were incubated for 30 min at 26°C in artificial seawater containing a final concentration of approximately 0.10 M $MgCl_2$ in the annular tracking chamber to suppress active swimming [2]. Larvae were classified as chemically immobilized when no active swimming or reorientation was observed and their subsequent motion consisted of an approximately straight, constant-speed vertical rise. For both developmental stages, we selected a common time interval from the vertical trajectory and corresponding image sequence. A linear fit to $z(t)$ provided the mean terminal rise speed and verified that motion was approximately constant over the selected interval (Fig. S1a). To obtain a frame-resolved velocity, we additionally fit $z(t)$ with a cubic smoothing spline and evaluated its derivative at each image time. We define

$$U(t) = \frac{dz}{dt}, \tag{S1}$$

so that an upward-rising, positively buoyant larva has $U > 0$. In each corresponding image (Fig. S1b), the larval silhouette was segmented and represented by its equivalent ellipse. The associated prolate spheroid had semi-major axis $a$ and equal transverse semi-axes $b$, giving

$$V_{\text{ell}}(t) = \frac{4\pi}{3} a(t) b^2(t) \tag{S2}$$

as an estimate of its volume. Fig. S1c shows the 3D prolate spheroid fit from the measured values of $a$ and $b$. Because the larvae remained approximately aligned with the direction of rise over the selected intervals, we used the Stokes resistance for translation parallel to the spheroid major axis. Writing $\zeta_\parallel = \eta D_\parallel$, the resistance length is

$$D_\parallel = \frac{16\pi L}{(s^2+1)\ln\left(\frac{s+1}{s-1}\right) - 2s}, \tag{S3}$$

where

$$L = (a^2 - b^2)^{1/2}, \qquad E = \frac{a}{b}, \qquad s = \frac{E}{(E^2-1)^{1/2}}. \tag{S4}$$

Here, $\eta$ is the dynamic viscosity of the medium in which the trajectory was recorded. For measurements performed in artificial seawater after treatment and rinsing, we used $\eta = 0.97 \times 10^{-3}$ Pa s. Defining the buoyant density deficit as

$$\Delta\rho = \rho_{\text{water}} - \rho_{\text{body}}, \tag{S5}$$

the terminal force balance gives

$$\zeta_\parallel(t) U(t) = \Delta\rho(t) V_{\text{ell}}(t) g, \qquad \Delta\rho(t) = \frac{\eta D_\parallel(t) U(t)}{V_{\text{ell}}(t) g}, \tag{S6}$$

where $g = 9.81$ m s$^{-2}$. Thus, upward motion gives $\Delta\rho > 0$, corresponding to a body that is less dense than the surrounding fluid (Fig. S1d). The density estimates are summarized in Fig. S1e. Along with the density estimate at 1 dpf, these measurements provide the two developmental anchors used in Fig. 4c(iv). Because equivalent passive measurements were not performed at every intermediate developmental stage, $\Delta\rho$ was linearly interpolated between the dpf 1 and dpf 9 anchor values.

The mean density deficit was 8.7 kg m$^{-3}$ at 1 dpf (range 4.1–14 kg m$^{-3}$) and 2.4 kg m$^{-3}$ at 9 dpf (range 1.0–3.2 kg m$^{-3}$). We then linearly interpolated the central estimate and both bounds between these two ages to estimate values at stages without passive-rise measurements. At dpf 0–1, active propulsion was set to zero and the measured terminal rise velocity was used directly as the buoyant drift. Published lipid measurements provide context for the developmental decline in buoyancy [3–5].

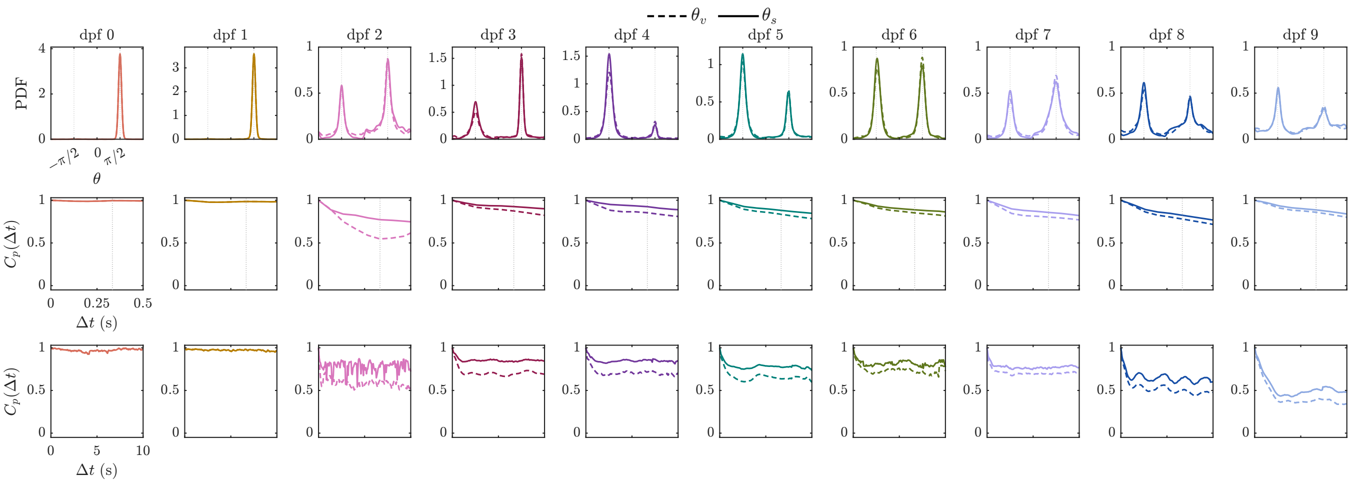


FIG. S2. **Effect of buoyancy correction on heading statistics.** Comparison of the trajectory heading $\theta_v = \mathrm{atan2}(\dot{z}, \dot{x})$ (dashed) and the inferred swimming heading $\theta_s = \mathrm{atan2}(\dot{z} - v_b, \dot{x})$ (solid), where $v_b$ is the estimated track-specific buoyant drift. Columns show successive developmental stages, with color indicating dpf. The top row shows heading probability densities; the middle and bottom rows show the corresponding heading autocorrelations over 0–0.5 s and 0–10 s, respectively. Statistics combine the upward and downward modes according to their track fractions, with equal track weighting within each mode.

#### C. Separation of buoyant drift and active swimming speed

For actively swimming larvae, we used the density deficits estimated above to calculate the buoyant contribution to each trajectory. Within each dpf, trajectories were first grouped into upward and downward swimming modes according to the sign of their net vertical displacement, $z(t_{\mathrm{end}}) - z(t_{\mathrm{start}})$.

The two-dimensional model (Eq. 3 of the main text) separates the buoyant contribution from active propulsion:

$$v_x(t) = v_s \cos\theta_s(t), \tag{S7}$$

$$v_z(t) = v_b + v_s \sin\theta_s(t). \tag{S8}$$

Here, positive $v_b$ denotes upward buoyant rise, and $v_s \geq 0$ is the active swimming speed along the inferred heading $\theta_s$. For track $j$ at a specific developmental stage (dpf), we combined the interpolated density deficit $\Delta\rho$ with the volume and resistance calculated from each image. Rearranging the terminal force balance from the preceding section gives

$$v_{b,j}(t) = \frac{\Delta\rho\, g\, V_{\mathrm{ell},j}(t)}{\zeta_{\parallel,j}(t)}, \qquad \zeta_{\parallel,j}(t) = \eta D_{\parallel,j}(t). \tag{S9}$$

The use of the parallel spheroidal resistance assumes that the major body axis is approximately aligned with the direction of passive translation. We assigned a single buoyant velocity to each track by averaging these estimates over time:

$$v_{b,j} = \langle v_{b,j}(t) \rangle_{w,j}\,, \qquad w_{ij} = \frac{\Delta t_{ij}}{\sum_k \Delta t_{kj}}, \tag{S10}$$

where $\Delta t_{ij}$ is the time interval represented by sample $i$ in track $j$.

For actively swimming stages, we subtracted this track-level buoyant drift before estimating the swimming heading, $\theta_{s,j}(t) = \mathrm{atan2}[v_{z,j}(t) - v_{b,j}, v_{x,j}(t)]$. This differs from the trajectory heading $\theta_v = \mathrm{atan2}(v_z, v_x)$ used in Fig. 3. At 0–1 dpf, where $v_s = 0$, $\theta_v$ describes passive motion and is not interpreted as an active swimming direction. Figure S2 compares the heading probability densities and autocorrelations before and after buoyancy correction across development.

With $v_{b,j}$ fixed, the active speed was obtained by minimizing the weighted residual of both measured velocity components:

$$\mathcal{L}_j(v_s) = \sum_i w_{ij} \left[ (v_{x,ij} - v_s \cos\theta_{s,ij})^2 + (v_{z,ij} - v_{b,j} - v_s \sin\theta_{s,ij})^2 \right]. \tag{S11}$$

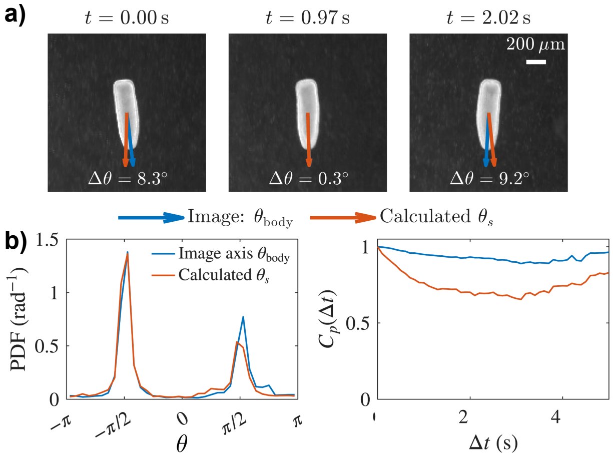


FIG. S3. **Comparison of image-derived body orientation and buoyancy-corrected swimming heading.** Data were obtained from nine trials at 5 dpf, the midpoint of the linear density interpolation between the measurements at 1 and 9 dpf. (a) Sequential snapshots from a representative recording, comparing the projected body-axis heading measured from the images, $\theta_{\rm body}$ (blue), with the inferred swimming heading, $\theta_s = {\rm atan2}(v_z - v_b, v_x)$ (orange). Here, $v_x$ and $v_z$ are the measured velocity components, and $v_b$ is the buoyant drift estimated from the interpolated density and measured body geometry. The image-based estimate assumes that propulsion is directed along the larval long axis, an alignment that need not hold in general. The absolute angular difference between the two headings is indicated by $\Delta\theta$. Times are relative to the first displayed frame. Scale bar, 200 $\mu$m. (b) Heading probability densities (left) and temporal heading autocorrelations, $C_p(\Delta t) = \langle \cos[\theta(t+\Delta t) - \theta(t)] \rangle$ (right), comparing $\theta_{\rm body}$ and $\theta_s$ across the nine trials using the same color convention.

The resulting least-squares estimator is

$$v_{s,j} = \frac{\langle v_x \cos\theta_s + (v_z - v_{b,j}) \sin\theta_s \rangle_{w,j}}{\langle \cos^2\theta_s + \sin^2\theta_s \rangle_{w,j}}, \tag{S12}$$

where the denominator is unity for an exactly normalized heading vector.

The resulting track-level $v_b$ and $v_s$ estimates were averaged with equal track weight separately for each dpf and swimming mode to obtain the values in Fig. 4c(v–vi). The difference in $v_s$ between the two modes was used to identify gravikinesis.

To propagate uncertainty in the density deficit, we repeated Eqs. (S9)–(S12) and the subsequent averaging using the lower and upper density bounds, $\Delta\rho^{(-)}$ and $\Delta\rho^{(+)}$. The error bars in Fig. 4c(v–vi) show the resulting extrema relative to the central estimate. These bounds reflect the experimental density ranges under the assumed linear interpolation between 1 and 9 dpf. They do not account for uncertainty associated with three-dimensional body tilt or departures from the equivalent prolate-spheroid approximation.

We additionally compared the inferred swimming heading $\theta_s$ with the projected body-axis heading $\theta_{\rm body}$ measured from images in nine trials at 5 dpf, the midpoint of the density interpolation between 1 and 9 dpf (Fig. S3). The image-based estimate treats the larval long axis as a proxy for the propulsion direction, although this alignment need not hold in general. Body curvature can complicate the definition of a single long axis, and propulsion may be oblique to that axis, therefore trajectory kinematics provides a more robust measure of propulsion direction than 2D body orientation.

### D. Fixation and immunostaining

To relate swimming behavior to ciliary and neural development, approximately 5–10 larvae were fixed almost daily from fertilization through approximately one month post fertilization, following Yuan et al. [6]. Larvae were transferred, with minimal seawater carryover, into microcentrifuge tubes containing 4% formaldehyde in 0.2 $\mu$m-filtered seawater. They were fixed for 1 h at room temperature in the dark and then washed twice in PBS. Larvae not stained immediately were dehydrated through a graded ethanol series (30%, 50%, 80% and 100%) and stored in 100% methanol at −20 °C. Larvae from 1, 5 and 12 dpf are shown in Fig. 2c.

Antibody labeling also followed Yuan et al. [6]. Stored larvae were rehydrated through the ethanol series and washed for 2 h in PBSTr (PBS + 0.3% Triton X-100). They were then blocked for 1 h at room temperature in 3%

bovine serum albumin (BSA) in PBSTr. Thirty larvae were incubated for 24 h at 4 °C in 4 mL of PBSTr containing two primary antibodies: rabbit polyclonal anti-FMRFamide antiserum (1:160; Chemicon/MilliporeSigma, AB15348) and mouse monoclonal anti-acetylated $\alpha$-tubulin (1:800; clone 6-11B-1, Sigma-Aldrich, T7451). Larvae were then washed in PBSTr for 2 h and blocked again in 3% BSA in PBSTr for 1 h at room temperature. Next, they were incubated overnight at 4 °C with two secondary antibodies, both at 1:100: goat anti-mouse IgG (H+L) Alexa Fluor 488, cross-adsorbed (Invitrogen, A-11001), and goat anti-rabbit IgG (H+L) Alexa Fluor 647, preadsorbed (Abcam, ab150087). After a final 2 h wash in PBSTr at room temperature, larvae were mounted in Vectashield and imaged on an inverted laser scanning confocal microscope (LSM 780/FLIM, Zeiss) equipped with a PLAN APO 63x/1.46 Oil objective lens (Zeiss). Image processing is conducted using both Zen software and Fiji (ImageJ). Images in Fig. 2c are maximum-intensity projections.

### III. THEORETICAL MODEL

#### A. Minimal projected swimmer model

##### 1. *Rotational diffusivity from the heading autocorrelation*

The estimator in Eq. 6 of the main text follows from the short-time behavior of the angular dynamics. Within each swimming mode, the preferred direction $\psi$ is fixed, and the aligned heading is $\eta = \theta_s - \psi$, wrapped to $[-\pi, \pi)$. The swimming-heading autocorrelation can therefore be written as

$$C_{p,s}(\Delta t) = \langle \cos\left[\eta(t+\Delta t) - \eta(t)\right] \rangle \,. \tag{S13}$$

Applying Itô's formula [7] to the orientation dynamics in Eq. 5 of the main text, with constant rotational diffusivity $D_r$, gives

$$\frac{\mathrm{d}C_{p,s}}{\mathrm{d}\Delta t} = -\left\langle a(\eta(t+\Delta t)) \sin\left[\eta(t+\Delta t) - \eta(t)\right] \right\rangle - D_r C_{p,s}(\Delta t). \tag{S14}$$

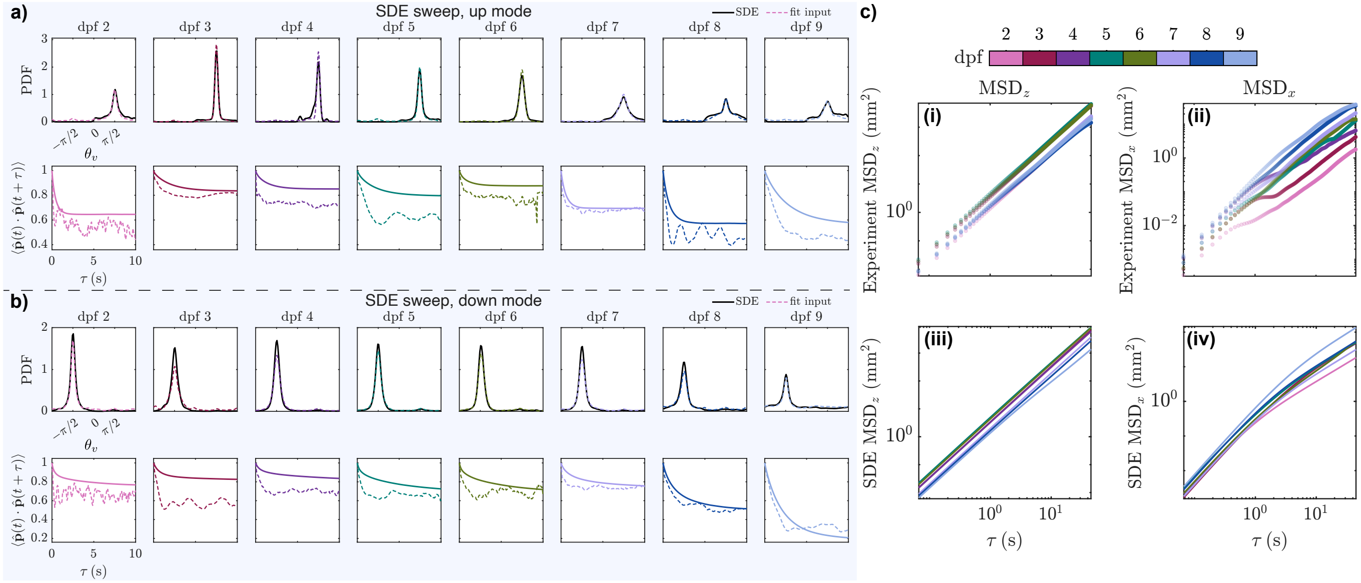


FIG. S4. **Comparison between experimental statistics and the minimal projected swimmer model.** (a,b) Stationary projected-heading distributions $\pi(\theta_v)$ (top rows) and projected-heading autocorrelations $C_p(\Delta t) = \langle \hat{\mathbf{p}}(t) \cdot \hat{\mathbf{p}}(t+\Delta t) \rangle$ (bottom rows) for the upward (a) and downward (b) swimming modes. Here, $\hat{\mathbf{p}} = (\cos\theta_v, \sin\theta_v)$ is the unit vector along the total projected velocity, including buoyancy. Dashed curves show experimental measurements and solid curves show simulations of Eqs. 3 and 5 of the main text using the stage- and mode-specific fits to $\pi(\eta)$ and $D_r$. Panels (a,b) show dpf 2–9. (c) Raw vertical and horizontal mean-squared displacements in the experiment (i,ii) and minimal-model simulations (iii,iv), with upward and downward trajectories pooled and color indicating dpf. The model reproduces the approximately ballistic vertical and diffusive horizontal scaling, supporting the use of the fitted orientation dynamics as a reduced description of dispersal-relevant behavior. The model coarse-grains, rather than phase-resolves, individual helical swimming cycles.

At zero lag, the sine term vanishes and $C_{p,s}(0) = 1$. The restoring drift therefore makes no contribution to the initial slope, giving

$$C_{p,s}(\Delta t) = 1 - D_r \Delta t + O(\Delta t^2), \tag{S15}$$

and hence

$$D_r = -\left.\frac{\mathrm{d}C_{p,s}}{\mathrm{d}\Delta t}\right|_{\Delta t = 0^+}. \tag{S16}$$

This result does not require a linear restoring drift or an exponential autocorrelation. The initial slope isolates the rotational diffusivity, whereas the correlation at longer lags also depends on the restoring dynamics. In practice, the derivative at zero lag was approximated by fitting the same short-lag portion of the mode-resolved correlation for every developmental stage.

##### 2. *Numerical validation of the minimal model*

For each developmental stage and swimming mode, numerical trajectories were generated from Eqs. 3 and 5 of the main text using the estimated values of $v_s$, $v_b$, and $D_r$ and the complete data-inferred drift $a(\eta)$. Initial aligned headings were sampled from the fitted stationary distribution $\pi(\eta)$, and $\theta_s = \eta + \psi$ was wrapped periodically. To compare with the experimental trajectory headings, we reconstructed $\theta_v = \mathrm{atan2}(v_b + v_s \sin\theta_s, v_s \cos\theta_s)$. Heading distributions, autocorrelations, and MSDs were then calculated using the same estimators as for the experimental data (Fig. S4).

Because the comparison uses $\theta_v$, the simulated heading distributions depend on the estimated buoyant drift and swimming speed as well as the fitted distribution of $\theta_s$. Their agreement provides a consistency check on these inputs. The autocorrelations beyond the short-lag interval used to estimate $D_r$, together with the MSDs, test whether the inferred angular dynamics reproduce motion over longer times. The simulations recover approximately ballistic vertical MSDs and approximately diffusive horizontal MSDs, although differences remain in the decay and long-lag values of the heading autocorrelations. These comparisons support the model as a description of the broad transport scaling, while the phase of individual helical cycles and the associated oscillations are not explicitly resolved.

#### B. Spectral construction and normalization of the Gaussian random flow

The background flow used in Fig. 5 and Fig. S5 is a zero-mean, incompressible Gaussian random field in the $x$–$z$ plane [8]. Its parameters are the rms speed $u_0 = \langle |\mathbf{u}|^2 \rangle^{1/2}$, spatial correlation scale $\ell_0$, and Eulerian correlation time $\tau_0$. Below, we describe its spectral construction.

The flow is generated on a periodic square of side $L$ from a Gaussian streamfunction,

$$\Psi(\mathbf{x}, t) = \sum_{\mathbf{k} \neq \mathbf{0}} B(k)\, a_{\mathbf{k}}(t) e^{i\mathbf{k}\cdot\mathbf{x}}, \qquad \mathbf{u} = \nabla^{\perp}\Psi = (\partial_z \Psi, -\partial_x \Psi), \tag{S17}$$

where $k = |\mathbf{k}|$. The Hermitian condition $a_{-\mathbf{k}} = a^*_{\mathbf{k}}$ ensures that $\Psi$ is real. The zero mode is removed, and the modal amplitudes have zero mean. The Gaussian spectral envelope is

$$B(k) \propto \exp\left(-\frac{k^2 \ell_0^2}{4}\right), \qquad |B(k)|^2 \propto \exp\left(-\frac{k^2 \ell_0^2}{2}\right), \tag{S18}$$

with the normalization chosen so that $\langle |\mathbf{u}|^2 \rangle = u_0^2$. Numerically, $\mathbf{k} = 2\pi(n_x, n_z)/L$, with $n_x, n_z = -M, \ldots, M$ and $M = 16$.

One member of each Hermitian pair evolves as an independent Ornstein–Uhlenbeck process. Over a flow time step $\Delta t$, the modal update is evaluated exactly as

$$a_{\mathbf{k}}(t + \Delta t) = e^{-\Delta t/\tau_0} a_{\mathbf{k}}(t) + \sqrt{1 - e^{-2\Delta t/\tau_0}}\, \xi_{\mathbf{k}}, \tag{S19}$$

where $\xi_{\mathbf{k}}$ is freshly sampled, zero-mean, unit-variance complex Gaussian noise with the same Hermitian symmetry. Each mode therefore has the temporal correlation $\langle a_{\mathbf{k}}(t) a^*_{\mathbf{k}}(t+s) \rangle \propto e^{-|s|/\tau_0}$. Thus, $\tau_0$ describes the temporal correlation of the flow at a fixed spatial position.

To relate $u_0$ and $\ell_0$ to the velocity-gradient magnitude, let $\mathbf{A} = \nabla\mathbf{u}$, with $\mathbf{A}:\mathbf{A} = \sum_{i,j} A_{ij}^2$. We define the dimensionless spectral coefficient $C_A$ through $\langle \mathbf{A}:\mathbf{A}\rangle = C_A u_0^2/\ell_0^2$. Its value is determined by the chosen spectrum. Since one spatial derivative of $\Psi$ produces $\mathbf{u}$, whereas two produce $\mathbf{A}$, the velocity and velocity-gradient variances involve the second and fourth spectral moments, respectively. In the continuum spectral limit, their ratio gives

$$C_A = \ell_0^2 \frac{\displaystyle\int_0^\infty k^5 e^{-k^2\ell_0^2/2}\,\mathrm{d}k}{\displaystyle\int_0^\infty k^3 e^{-k^2\ell_0^2/2}\,\mathrm{d}k} = 4. \tag{S20}$$

The reference dissipation rate $\varepsilon$ and kinematic viscosity $\nu$ define the Kolmogorov length, time, and velocity scales,

$$\eta_K = \left(\frac{\nu^3}{\varepsilon}\right)^{1/4}, \qquad \tau_K = \left(\frac{\nu}{\varepsilon}\right)^{1/2}, \qquad u_K = \frac{\eta_K}{\tau_K} = (\nu\varepsilon)^{1/4}. \tag{S21}$$

The matching condition in Eq. 9 of the main text therefore becomes

$$G_K = \langle \mathbf{A}:\mathbf{A}\rangle^{1/2} = \frac{2u_0}{\ell_0} = \left(\frac{\varepsilon}{\nu}\right)^{1/2} = \frac{1}{\tau_K}. \tag{S22}$$

This calibration is similar to the statistical-model matching approach reviewed by Bec et al. [8], which relates the velocity-gradient timescale to the Kolmogorov time while specifying the spatial scale and Eulerian persistence separately. We specify the spatial scale through $\ell_0 = a_\eta \eta_K$ and the Eulerian correlation time through the Kubo number $\mathrm{Ku} = u_0\tau_0/\ell_0$. The gradient-matching condition then gives

$$u_0 = \frac{a_\eta}{2} u_K, \qquad \tau_0 = \mathrm{Ku}\frac{\ell_0}{u_0} = 2\,\mathrm{Ku}\,\tau_K. \tag{S23}$$

The values $a_\eta = \mathrm{Ku} = 10$ used here therefore correspond to $\ell_0 = 10\eta_K$, $u_0 = 5u_K$, and $\tau_0 = 20\tau_K$. In the model, the Eulerian field consequently evolves slowly compared with the matched gradient timescale.

For completeness, the same spectral moments give the planar vorticity–strain relations

$$\langle \omega^2\rangle = \langle \mathbf{A}:\mathbf{A}\rangle, \qquad \langle \mathbf{E}:\mathbf{E}\rangle = \frac{1}{2}\langle \mathbf{A}:\mathbf{A}\rangle, \tag{S24}$$

where $\omega = \partial_x u_z - \partial_z u_x$ and $\mathbf{E} = (\mathbf{A} + \mathbf{A}^\top)/2$. These are relations within the two-dimensional GRF and don't imply equality of its individual gradient components with those of three-dimensional turbulence. Because $C_A = 4$ is fixed by the Gaussian spectrum, Fig. S5 tests sensitivity to the remaining choices, $a_\eta$ and Ku, about the values used in Fig. 5.

#### C. Effective heading-relaxation rate

To compare still-water heading dynamics with reorientation by flow, we summarize each swimming mode by an effective relaxation rate $\Omega_\theta$. This rate is calculated from the parameters $D_r$ and $A_1$. Defining the swimming-heading unit vector as $\hat{\mathbf{p}}_s = (\cos\theta_s, \sin\theta_s)$, the heading autocorrelation can be written equivalently as

$$\begin{aligned} C_{p,s}(\Delta t) &= \langle \hat{\mathbf{p}}_s(t)\cdot\hat{\mathbf{p}}_s(t+\Delta t)\rangle \\ &= \langle \cos\left[\eta(t+\Delta t) - \eta(t)\right]\rangle, \end{aligned} \tag{S25}$$

because the preferred direction $\psi$ in $\theta_s = \eta + \psi$ is fixed within each mode. Within a stationary swimming mode, headings at sufficiently long time separations become statistically independent while retaining the same directional bias. The unit-vector expression then gives

$$\lim_{\Delta t\to\infty} C_{p,s}(\Delta t) = \langle \hat{\mathbf{p}}_s\rangle \cdot \langle \hat{\mathbf{p}}_s\rangle = \left|\langle \hat{\mathbf{p}}_s\rangle\right|^2.$$

Equivalently, expanding the cosine of the angle difference and using independence gives

$$\begin{aligned} \lim_{\Delta t\to\infty} C_{p,s}(\Delta t) &= \langle \cos\eta\rangle^2 + \langle \sin\eta\rangle^2 \\ &= \left|\langle e^{i\eta}\rangle\right|^2 = A_1^2. \end{aligned} \tag{S26}$$

Thus, $A_1$ is also the length of the mean heading vector. Subtracting the fixed preferred angle $\psi$ rotates this vector without changing its length. Using the same exponential form as the fit defining $\tau_h$ in Eq. 2 of the main text, we approximate the correlation within each mode as

$$C_{p,s}(\Delta t) \simeq A_1^2 + \left(1 - A_1^2\right) e^{-\Omega_\theta \Delta t}. \tag{S27}$$

Here, $1 - A_1^2$ is the amount of correlation that decays, and $\Omega_\theta$ is its decay rate. The initial slope is $-(1 - A_1^2)\Omega_\theta$. Matching this to the slope $-D_r$ derived in Eq. (S15) gives

$$\Omega_\theta = \frac{D_r}{1 - A_1^2}. \tag{S28}$$

Thus, $\Omega_\theta^{-1}$ is the relaxation time of an exponential approximation that matches both the initial decay and the long-time correlation.

For the same exactly exponential correlation, this construction and the fit in Eq. 2 of the main text would give $C_\infty = A_1^2$ and $\Omega_\theta = 1/\tau_h$. The quantities reported here differ in both the heading variable and the grouping of trajectories. In Fig. 3d, $\tau_h$ is fitted to the trajectory-heading correlation for $\theta_v$, which includes buoyant motion and pools the upward and downward modes. Here, $\Omega_\theta$ is calculated from the buoyancy-corrected swimming heading $\theta_s$ separately within each mode.

The upward- and downward-mode rates were calculated separately. For each developmental stage, the value shown in the inset to Fig. 5d was their geometric mean,

$$\Omega_{\theta,\text{stage}} = \sqrt{\Omega_{\theta,\uparrow}\Omega_{\theta,\downarrow}}. \tag{S29}$$

This averaging also need not give the inverse of a relaxation time fitted to the pooled correlation. We compared $\Omega_{\theta,\text{stage}}$ with the imposed velocity-gradient rate at the dissipation rate $\varepsilon_c^{\text{ret}}$ where the simulations reached half retention,

$$G_{K,c} = \left(\frac{\varepsilon_c^{\text{ret}}}{\nu}\right)^{1/2}. \tag{S30}$$

More generally, balancing the flow-gradient rate with the still-water heading-relaxation rate gives

$$\varepsilon_\theta \sim \nu\Omega_\theta^2. \tag{S31}$$

This provides an order-of-magnitude estimate of the flow strength at which imposed rotation competes with still-water heading relaxation. The actual half-retention point is determined from the simulations.

#### D. Numerical implementation of transport simulations

For each developmental stage, swimming mode, and imposed flow strength in Fig. 5, we simulated 1,000 non-interacting particles in each of 32 independent flow realizations. We considered 25 logarithmically spaced values of $\varepsilon$ from $10^{-10}$ to $10^{-2}$ W kg$^{-1}$. The flow was defined on a periodic square of side $L = 20\ell_0$, ranging from 2 m at $\varepsilon = 10^{-10}$ W kg$^{-1}$ to 20 mm at $\varepsilon = 10^{-2}$ W kg$^{-1}$. Initial particle positions were distributed uniformly over this domain, and initial aligned headings were sampled from the fitted stationary distribution for the corresponding mode. Particle coordinates were reduced modulo $L$ when evaluating the periodic flow, while unwrapped positions were retained for displacement and transport calculations. We discarded a 20-s initialization period and retained the following 300 s, recording positions and headings every 1/15 s.

Active-swimmer dynamics were integrated using Euler–Maruyama with a time step of 1/6000 s. The fitted heading drift was evaluated by periodic linear interpolation, and headings were wrapped to $[-\pi, \pi)$ after each update. Flow coefficients were advanced using the exact Ornstein–Uhlenbeck update described above. The flow-update frequency increased with $\varepsilon$, targeting update intervals of approximately $0.02\tau_K$ or shorter, where $\tau_K = (\nu/\varepsilon)^{1/2}$. Flow velocities and gradients were evaluated at the current particle positions at each flow update and held fixed between updates. Nonpropelling particles used integration steps of 1/750 s for the 20 lowest dissipation rates and 1/6000 s for the five highest.

The effective vertical velocity was obtained from the mean net vertical displacement, and $D_x$ was estimated as half the slope of a linear fit to the horizontal MSD over $5 < \Delta t \leq 60$ s. A common neutrally buoyant passive-tracer baseline, with $v_s = v_b = 0$, was used at each dissipation rate. Standard errors were estimated across the 32 flow realizations and propagated to the normalized transport metrics, holding the fitted biological parameters fixed.

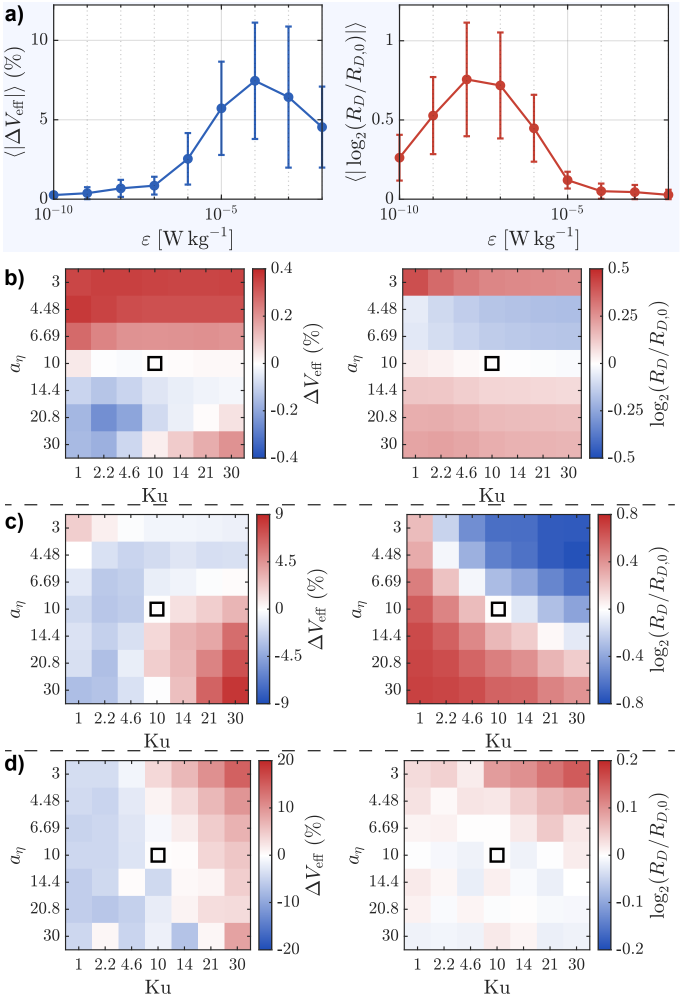


FIG. S5. **Sensitivity of the Figure 5 transport predictions to the Gaussian-random-flow closure parameters.** The closure grid varies the spatial-scale factor $a_\eta$, defined by $\ell_0 = a_\eta \eta_K$, and the Kubo number $\mathrm{Ku} = u_0\tau_0/\ell_0$ about the final choice $(a_\eta, \mathrm{Ku}) = (10, 10)$ which gives $V_{\mathrm{eff},0}$) and $R_{D,0}$. (a) Mean closure sensitivity across the full dissipation-rate sweep for the effective vertical velocity $V_{\mathrm{eff}}$ (left) and the normalized horizontal diffusivity $R_D = D_x^{\mathrm{active}}/D_x^{\mathrm{passive}}$ (right). For each developmental stage and closure-grid cell, the deviations are calculated as: $\Delta V_{\mathrm{eff}} = 100(V_{\mathrm{eff}} - V_{\mathrm{eff},0})/|V_{\mathrm{eff},0}|$ and $\log_2(R_D/R_{D,0})$. The markers denote the mean over dpf 3, 6, and 9 and as $a_\eta$ and Ku are varied. The vertical error bars denote $\pm 1$ standard deviation across those cells. (b–d) Signed, dpf-averaged closure sensitivities at increasing reference dissipation rates, $\varepsilon = 10^{-10}$, $10^{-6}$, and $10^{-2}\,\mathrm{W\,kg^{-1}}$, respectively. Within each panel, the left heat map shows $\Delta V_{\mathrm{eff}}$, while the right heat map shows $\log_2(R_D/R_{D,0})$. Red and blue indicate increases and decreases, respectively, relative to the production closure, which is marked by the black square.

#### E. Sensitivity to the GRF closure parameters

The Gaussian spectral kernel fixes $C_A = 4$, leaving the spatial-scale factor $a_\eta$ and the Kubo number Ku as the two dimensionless closure choices. To test the dependence of the transport predictions on these choices, the full $\varepsilon$ sweep (Fig. S5) was repeated over a $7 \times 7$ grid spanning

$$3 \le a_\eta \le 30, \qquad 1 \le \mathrm{Ku} \le 30, \tag{S32}$$

while Fig. 5 in the main text uses

$$(a_\eta, \mathrm{Ku}) = (10, 10). \tag{S33}$$

For these simulations, we used the normal up-swimming modes of dpf 3, 6, and 9, together with the common passive-tracer sweep used to define $R_D$. For each stage, dissipation rate, and closure-grid cell, the effective vertical-velocity deviation relative to that stage's production baseline was

$$\Delta V_{\mathrm{eff}} = 100\frac{V_{\mathrm{eff}} - V_{\mathrm{eff},0}}{|V_{\mathrm{eff},0}|}, \tag{S34}$$

and the diffusivity-ratio deviation was

$$\Delta_R = \log_2\left(\frac{R_D}{R_{D,0}}\right). \tag{S35}$$

In the sweep-averaged summary (Fig. S5a), $|\Delta V_{\mathrm{eff}}|$ and $|\Delta_R|$ were evaluated before averaging, while the signed heat maps retain the direction of the change. The variation of the swimming response is weak at the lowest $\varepsilon$, but becomes most sensitive over parts of the transition where active transport is being eroded, and decreases again once the swimmers approach their high-gradient limits. Figure S5 supports the qualitative robustness of the Fig. 5 transport trends while showing that the absolute magnitudes and precise crossover locations retain some dependence on the spatial and temporal closure scales.


[1] A. Takeda-Sakazume, J. Honjo, S. Sasano, K. Matsushima, S. A. Baba, Y. Mogami, and M. Hatta, Gravitactic Swimming of the Planula Larva of the Coral Acropora: Characterization of Straightforward Vertical Swimming, Zoological Science **40**, 44 (2022).

[2] Q. Zhang, E. Bollati, C. O. Pacherres, D. J. Hughes, W. Dellisanti, C. Ferrier-Pages, and M. Kühl, Active host control of the internal o2 microenvironment in reef-building corals, Iscience **29** (2026).

[3] J. Figueiredo, A. H. Baird, M. F. Cohen, J.-F. Flot, T. Kamiki, T. Meziane, M. Tsuchiya, and H. Yamasaki, Ontogenetic change in the lipid and fatty acid composition of scleractinian coral larvae, Coral Reefs **31**, 613 (2012).

[4] S. Harii, K. Nadaoka, M. Yamamoto, and K. Iwao, Temporal changes in settlement, lipid content and lipid composition of larvae of the spawning hermatypic coral Acropora tenuis, Marine Ecology Progress Series **346**, 89 (2007).

[5] I. Arai, M. Kato, A. Heyward, Y. Ikeda, T. Iizuka, and T. Maruyama, Lipid composition of positively buoyant eggs of reef building corals, Coral Reefs **12**, 71 (1993).

[6] D. Yuan, N. Nakanishi, D. K. Jacobs, and V. Hartenstein, Embryonic development and metamorphosis of the scyphozoan aurelia, Development genes and evolution **218**, 525 (2008).

[7] B. Oksendal, *Stochastic differential equations: an introduction with applications* (Springer Science & Business Media, 2013).

[8] J. Bec, K. Gustavsson, and B. Mehlig, Statistical models for the dynamics of heavy particles in turbulence, Annual Review of Fluid Mechanics **56**, 189 (2024).
